# Benchmarking Real-World Applicability of Molecular Generative Models from De novo Design to Lead Optimization with MolGenBench

**DOI:** 10.1101/2025.11.03.686215

**Authors:** Duanhua Cao, Zhehuan Fan, Jie Yu, Mingan Chen, Xinyu Jiang, Xia Sheng, Xingyou Wang, Chuanlong Zeng, Xiaomin Luo, Dan Teng, Mingyue Zheng

## Abstract

Structure-based drug design (SBDD) has been profoundly reshaped by the advent of deep generative models, yet their practical impact on drug discovery remains limited. A central issue is the absence of a rigorous, application-oriented benchmark that mirrors the multi-stage, target-aware workflows of real-world pharmaceutical development. Inspired by recent advances in benchmarking for computer vision and large language models, where systematic evaluation has catalysed rapid progress, we introduce MolGenBench, a comprehensive benchmark designed to close the gap between molecular generation algorithms and tangible drug discovery outcomes. MolGenBench integrates a structurally diverse, large-scale dataset spanning 120 protein targets, 5,433 chemical series comprising 220,005 experimentally confirmed active molecules. Beyond conventional de novo generation, it incorporates a dedicated hit-to-lead (H2L) optimization scenario, which represents a critical phase in hit optimization that is seldom evaluated in existing benchmarks. We further introduce novel, pharmaceutically grounded metrics that assess a model’s ability to both rediscover target-specific actives or progressively optimize compounds for potency. Through extensive evaluation, MolGenBench uncovers significant gaps between current generative models and the demands of real-world drug development, establishing a foundational resource for building generative models with enhanced practical impact and accelerated translational potential.

## INTRODUCTION

The discovery of novel, safe, and effective therapeutics from chemical space to address the medical needs of billions worldwide and improve human quality of life remains a formidable challenge. Throughout recorded history, humanity has explored only a minuscule fraction of this vast chemical space, yet such exploration has yielded a wealth of molecules that underpin modern medical practice^1^. These successes achieved within limited exploration boundaries have inspired researchers to develop methods capable of efficiently identifying drug candidates across broader chemical landscapes, thereby addressing the escalating global burden of disease^2^. The convergence of artificial intelligence (AI) and life sciences is now driving transformative advances in this field. Recent years have witnessed significant progress in deep generative models for drug discovery^3–5^. Such approaches leverage target protein structural and functional information to propose novel drug-like molecules, offering a paradigm shift toward exhaustive exploration of unbounded chemical space while circumventing the constraints of conventional library-dependent screening^6^.

Deep generative models can design target-specific molecules by leveraging either protein sequences or structural information^7,8^. In recent years, the field has increasingly prioritized structure-based approaches. For instance, liGAN^8^ encodes binding pocket structures into atomic density grid representations, employing a conditional variational autoencoder (CVAE) to extract spatial features, predict molecular density grids, and generate molecules via rule-based atom fitting and bond inference algorithms. However, such methods suffer from inherent limitations in maintaining rotational and translational equivariance. To address this, subsequent models have adopted equivariant graph neural networks (E-GNNs) paired with autoregressive atom-by-atom generation frameworks^5,9–13^. Representative examples include SBDD^9^ and GraphBP^10^, which process structural data through E-GNNs to preserve geometric equivariance. Notably, these methods generate only atom types and coordinates, relying on external tools like RDKit^14^ for bond assignment. Recent advances further integrate end-to-end bond prediction. Pocket2Mol introduces a geometric vector perceptron (GVP)-based E-GNN to encode pocket context and directly predict edge types. This paradigm has inspired diverse innovations in pocket encoding and atomic sampling, including DeepICL^11^, SurfGen^12^, Delete^13^, and PocketFlow^5^.

Diverging from autoregressive approaches, TargetDiff^15^, PMDM^16^, DecompDiff^17^, and DiffSBDD^18^ employ diffusion models to generate complete molecular structures conditioned on protein pocket environments. These methods demonstrate distinct generative advantages through their unique molecular representations and loss function designs. Notably, MolCraft^19^ innovatively introduces Bayesian Flow Networks (BFN) to structure-based molecular generation (SBMG), significantly improving the three-dimensional conformational quality of generated molecules. With advancements in large language model (LLM) technology, researchers have begun exploring their application in SBMG. For instance, Lingo3DMol^20^ and 3DMolFormer^21^ utilize language models to jointly generate both atom types and spatial coordinates. Throughout this field’s development, various models have incorporated different prior knowledge to enhance generative capabilities: DecompDiff and Lingo3DMol employ fragment-based decomposition representations; SurfGen and Delete integrate protein surface physicochemical features; DeepICL and PGMG incorporate interaction information and pharmacophore constraints, respectively. The integration of these prior knowledge elements has effectively improved model performance in specific aspects.

Alongside de novo design, H2L optimization constitutes another critical task of drug design that focuses on improving compound properties (e.g., binding affinity and selectivity) through partial atomic modifications, typically by exploring R-group variations while maintaining core scaffolds. Compared to de novo molecule generation, this research area remains relatively underexplored. Autoregressive models demonstrate inherent suitability for this fragment-based design paradigm. Among diffusion-based approaches, DiffSBDD has shown promising results using inpainting techniques for fixed-fragment extensions ^18^. Notably, specialized methods like DiffDec^22^ have been developed specifically for this optimization scenario.

While generative models have advanced rapidly, methodologies and benchmark datasets for evaluation remain critically underdeveloped. Earlier benchmarking efforts systematically compared model performance but focused solely on ligand-centric properties without protein context^23,24^. Current practices predominantly rely on the CrossDocked2020^25^ dataset (hereafter referred to simply as CrossDock) which is an extended version of PDBBind^26^ for model training and testing. However, this dataset’s limitation of containing very few reference ligands per target fails to capture the complete distributional properties of active binding spaces, significantly compromising the real-world relevance of evaluation results^27^. Existing metrics often do not cover all relevant aspects of drug design projects. The AddCarbon model starkly illustrates this limitation by simply inserting random carbon atoms into training set molecules, it achieves near-perfect scores on standard metrics (uniqueness, diversity, novelty) while outperforming sophisticated generative models^28^. Furthermore, studies reveal strong correlations between some standard metrics (QED, Vina score) and atom counts, enabling even random ZINC^29^ database samples to outperform reference molecules in docking scores^30^. Existing metrics for evaluating synthesizability are similarly flawed. For instance, the Synthetic Accessibility score^31^ relies on the occurrence frequency of chemical substructures in curated databases to estimate synthetic feasibility, while the Synthetic Complexity score^32^ is trained on data from Reaxys^33^, a comprehensive reaction database to quantify molecular complexity, with an implicit encoding of the number of synthetic steps required. Both metrics along with others like the length of Simplified Molecular-Input Line-Entry System (SMILES) demonstrate poor discrimination between synthetically feasible and infeasible compounds when validated through retrosynthetic analysis^34^, further highlighting the need for more pharmacologically grounded synthesizability assessments in molecular generation benchmarks. Taken together, these findings highlight the urgent need for more rigorous, real-world relevance of evaluation metrics. Recent approaches now incorporate multidimensional assessment metrics that extend beyond basic chemical properties to include conformational quality, interaction patterns, and pharmacophore similarity. For example, PGMG introduced pharmacophore similarity metrics, while DeepICL developed interaction fingerprint assessments both designed to evaluate binding modes. Additionally, PoseCheck^35^ established 3D-specific evaluations, including strain energy, clash scores, and conformation similarity between generated and docked poses. However, these evaluations remain limited in scope, often tested on small target sets. For instance, PoseCheck benchmarked only five models using 3D metrics. Recent efforts have established new benchmarks for SBMG. Durian^27^ included fourteen target proteins with corresponding active compounds, enabling systematic comparison of six state-of-the-art models and POKMOL-3D^36^ assembled an even more comprehensive benchmark featuring thirty-two diverse targets with validated active compounds for evaluating nine generative approaches.

Although these efforts have made significant progress, enabling newly developed models to be evaluated more comprehensively on larger test sets, the current benchmarks still suffer from several limitations that hinder their ability to accurately reflect real-world applicability. These shortcomings can be categorized into four key areas:

i. **Dataset construction**: Overly stringent activity cutoffs in some datasets exclude experimentally validated active molecules, leading to systematic underestimation of models’ capacity to rediscover known bioactive compounds. Furthermore, compared to well-established virtual screening benchmarks (e.g., DUD-E^37^ and DEKOIS2.0^30^), the number and structural diversity of protein targets in existing molecular generation benchmarks remain inadequate to comprehensively assess a method’s broad utility in real-world scenarios.
ii. **Model selection**: The scope of evaluated model architectures is narrow, for example, BFN-based models are not included. Additionally, there is insufficient analysis of how variations in prior knowledge integration (e.g., pharmacophore constraints, protein surface features) and training data composition (e.g., simulated vs. crystal-derived protein-ligand complexes) influence model performance, limiting insights into design principles for improved methods.
iii. **Evaluation scenarios**: Current benchmarks exclusively focus on de novo molecule generation, while H2L optimization, such as fragment-guided molecular modification, a core step in advancing early-stage hits to drug-like leads, remains largexly unaddressed due to the lack of standardized assessment protocols.
iv. **Evaluation metrics**: Existing metrics fail to robustly evaluate two critical capabilities: first, a method’s capacity to leverage target-specific information (a prerequisite for generating target-specific molecules); second, the efficiency and efficacy of a model in identifying active molecules or scaffolds, an assessment where virtual screening benchmarks have long set a higher standard.

To address these challenges in evaluating molecular generation methods, we developed MolGenBench, a benchmark that better aligns with real-world drug discovery requirements. Firstly, we collected extensive experimentally validated active molecules across diverse targets to enable evaluation based on real bioactivity data rather than computational metrics. Secondly, we constructed congeneric series for these targets to assess lead optimization capabilities. Thirdly, we incorporated both conventional 2D/3D metrics for preliminary evaluation and newly designed metrics to comprehensively assess target-aware ability, active molecule generation capacity and efficiency and molecular optimization performance.

Leveraging our newly constructed MolGenBench dataset spanning 120 protein targets, 5,433 chemical series comprising 220,005 experimentally confirmed active molecules and tailored evaluation metrics (e.g., Target-Aware Score, MNA Score), we systematically tested molecular generation models encompassing diverse molecular representation schemes (2D vs. 3D), architectural paradigms (autoregressive, diffusion, Bayesian Flow Networks), integrated prior knowledge (pharmacophore constraints, protein surface features), and training data sources (simulated complexes like CrossDock vs. crystal-derived data like BindingMOAD). This design allowed us to disentangle the impact of each factor on model performance.

Our evaluation identifies three critical limitations of state-of-the-art approaches: First, both de novo and H2L models demonstrate limited ability to rediscover known active molecules at the molecular level, highlighting fundamental challenges in bioactive chemical space navigation, where precise steric and electronic complementarity to the target pocket is required. Second, even when generating chemically plausible molecules, notable deficiencies remain in predicting physically valid molecular conformations (e.g., high strain energy, steric clashes with the pocket), indicating inadequate integration of 3D structural constraints during model training or inference. Most critically, many structure-based models demonstrably fail to effectively leverage protein structural information during generation, despite this being their purported advantage, a disconnect that underscores the need for more powerful training paradigms that better encode protein-ligand interaction principles.

We contend that while deep generative models have advanced methodologically in molecular design (e.g., innovations in E-GNNs or diffusion frameworks), their limited practical impact in drug discovery stems largely from the absence of benchmarking standards and metrics that faithfully emulate real-world constraints (e.g., multi-stage optimization, novel target generalization). This study is a step towards closing this gap by establishing an assessment framework that faithfully recapitulates real-world requirements, thereby providing a foundation for advancing molecular generation technologies toward clinically meaningful applications.

## RESULTS AND DISCUSSION

### MolGenBench Dataset Construction and Baseline Model Selection

As shown in **Fig. 1a**, the data processing pipeline commenced with the acquisition of the ChEMBL^38^ database (version 33). The data underwent preprocessing, in which any ligand that failed RDKit’s molecular preparation step (due to invalid SMILES representation) was discarded. Affinity data labeled as NaN, 0, or containing symbols such as ‘<’ or ‘>’ were also excluded. Concurrently, data points with binding affinity values exceeding 10μM were excluded. This lenient threshold was intentionally set to maximize the retention of positive molecules and to ensure a diverse representation of experimentally confirmed actives across all protein targets. This approach mitigates the risk of underestimation in subsequent testing due to an insufficient number of positive reference compounds. During the subsequent clustering phase, the data were first partitioned by UniProt^39^ ID to group all entries associated with the same protein target. Within each UniProt ID group, affinity data were further organized into distinct Chemical Series based on ‘assay ID’ criteria. This grouping strategy ensured that all data within a given series were derived from identical assay conditions, with minimized systematic error in the binding affinity data to guarantee the accuracy of subsequent evaluations.

**Fig. 1.**
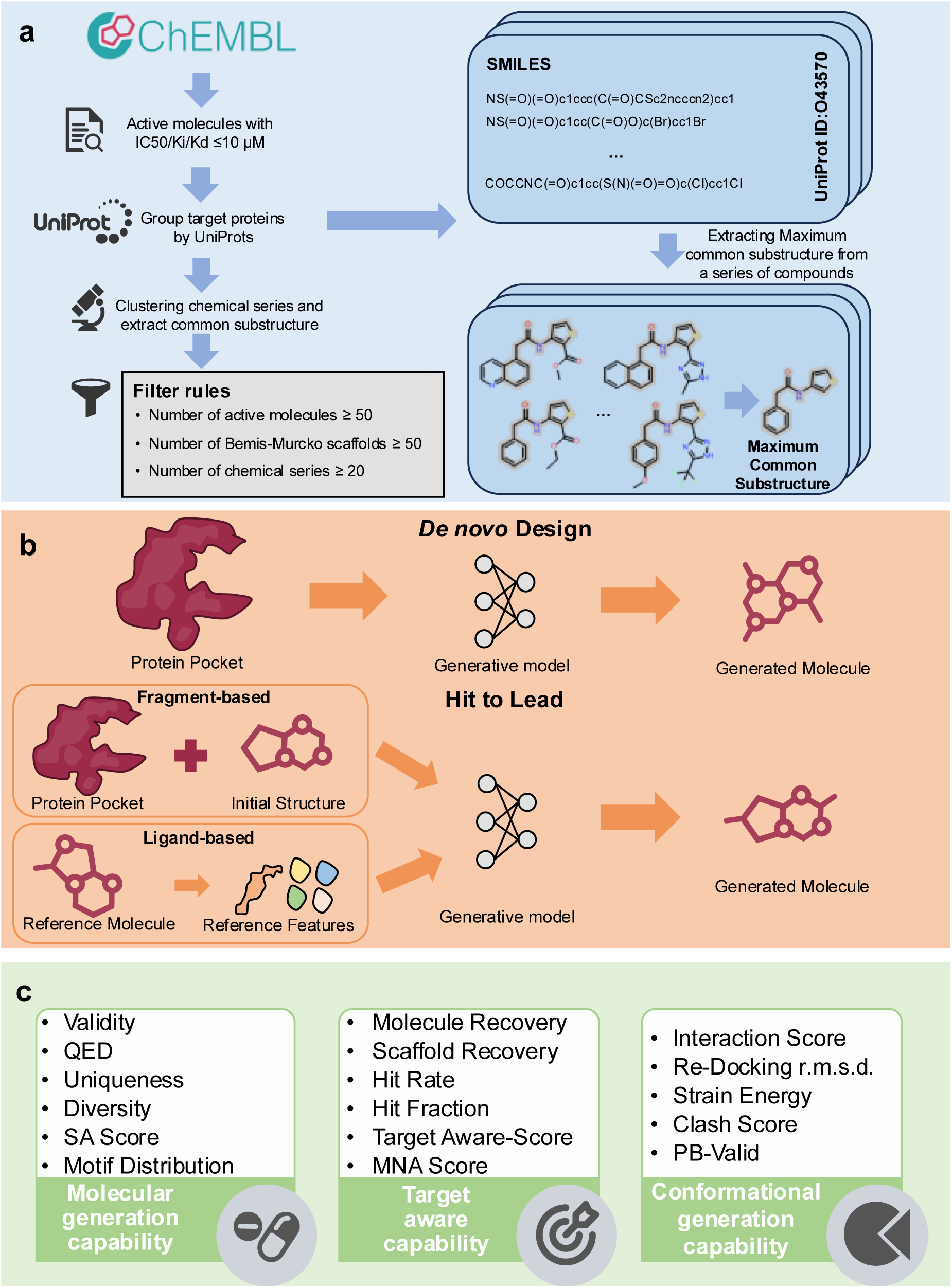
Core components of the MolGenBench benchmarking framework. **a,** Dataset collection and processing workflow. **b,** Dual evaluation scenarios aligned with drug discovery workflows: (i) *De novo molecular design*: Models generate molecules solely from the target protein information; (ii) *Hit-to-lead optimization*: Models generate molecules based on initial fragments (fragment-based) or reference actives features (ligand-based), mimicking the critical H2L stage rarely evaluated in existing benchmarks. **c,** Multi-dimensional evaluation metrics: Metrics address key drug discovery requirements, including: basic molecular properties, target-aware capability, and conformational quality.

For each UniProt ID, we quantified the number of unique SMILES strings, the count of distinct Bemis-Murcko scaffolds, and the number of chemical series. A subset with 120 UniProt IDs was then selected for the de novo molecular generation test set based on the following minimum thresholds: at least 50 active molecules, no fewer than 50 unique Bemis-Murcko scaffolds, and a minimum of 20 distinct chemical series. This stringent filtering guarantees that each tested protein system is supported by a substantial corpus of active compounds for reference, while also ensuring significant structural and chemical diversity among these actives. To construct a dedicated benchmark dataset for H2L optimization, we implemented an additional filtering and selection process for the compounds within each Series ID. Specifically, for a series to be considered, it was required to contain a common substructure. We identified the Maximum Common Substructure (MCS) for each series, applying a dual threshold: the MCS had to be present in over 80% of the molecules within the series, and its size had to constitute more than one-third of the atoms in any molecule containing it. Molecules failing to meet these criteria were discarded from subsequent analysis. Concurrently, we collected all available structures for the relevant targets that contained a co-crystallized ligand from the Protein Data Bank (PDB) database^40^. From these, a single PDB entry was selected for each target in subsequent molecular docking operations.

Finally, to build the final H2L optimization test set, we selected the top five series (ranked by the number of successfully docked ligands) within each UniProt ID. This resulted in a comprehensive dataset encompassing 120 unique protein targets and 600 distinct compound series, ensuring both structural and target diversity. Detailed protocols for the MCS analysis, ligand preparation, and docking procedures are provided in the Methods section. The detailed composition of our dataset is summarized in **SI-Fig. 1**.

To systematically disentangle the impacts of key methodological variables, including architectural paradigms, training data sources, and integrated prior knowledge, we selected a representative set of de novo generative models that collectively incorporate these variations, a critical design choice aligned with MolGenBench’s goal of addressing gaps in existing benchmarks. The models cover three core architectural families with distinct design logics: (1) autoregressive approaches (Pocket2Mol, leveraging GVP-based E-GNN networks; FLAG, integrating fragment-decomposed constraints; SurfGen, incorporating protein surface physicochemical features; TamGen, a target-aware chemical language model; and PocketFlow, incorporating explicit chemical rules), (2) diffusion models (TargetDiff, utilizing 3D equivariant diffusion; DecompDiff, adopting fragment-based decomposition priors; and DiffSBDD, which have two variants: DiffSBDD-M trained on BindingMOAD crystal complexes and DiffSBDD-C on CrossDock simulated complexes), and (3) Bayesian Flow Networks (BFNs), represented MolCraft, which optimizes 3D conformational quality via continuous parameter space modeling. Training data sources exhibited deliberate diversity to assess data-driven biases: large-scale molecular pre-training data (e.g., PubChem for TamGen, ZINC for PocketFlow) that prioritize broad chemical space coverage, and protein-ligand complex datasets (BindingMOAD, CrossDock) that focus on ligand-pocket interaction patterns. Furthermore, the models integrated different forms of structural and chemical priors, including fragment-based constraints (DecompDiff, FLAG) to guide generation via molecular decomposition, molecular surface information (SurfGen) to align ligand-pocket complementarity, and explicit chemical rules (PocketFlow) to enhance chemical feasibility. This diverse model panel enabled a comprehensive assessment of how various inductive biases shape generative performance, a central objective of MolGenBench’s benchmarking framework.

Building on our evaluation of de novo molecular generation models, we next focused on H2L optimization, a critical yet historically understudied stage in SBDD, by establishing a unified conceptual framework for its computational modeling. Herein, we frame generative models that commence from a pre-defined chemical structure, such as a complete ligand or a substructural fragment, as computational embodiments of the H2L optimization process. This perspective establishes a unified conceptual framework that encompasses both ligand-based and fragment-based approaches, defined by their exploitation of an initial chemical starting point. This stands in fundamental contrast to de novo design, which is initiated purely from a protein pocket. The “H2L” descriptor, therefore, offers a precise and efficient communicative shorthand for the scope and intent of these constraint-driven generative models.

In our assessment of H2L models, we included both fragment-based (DiffSBDD, Delete, DiffDec) and ligand-based (ShEPhERD, ShapeMol, PGMG) approaches. Fragment-based methods were provided with an initial fragment and its binding conformation in pocket, while ligand-based methods used features derived from reference active molecules as conditional inputs. The DiffDec model is uniquely characterized by its requirement for a scaffold SMILES string with pre-specified atomic anchors for modification (detailed procedures are described in the **Supplementary Information**). Architecturally, the models cover autoregressive frameworks (Delete, PGMG) and diffusion-based generative schemes (DiffDec, DiffSBDD). Training data varied across models and included specialized molecular pre-training datasets such as ChEMBL (PGMG) and MOSES^41^ (ShapeMol and ShEPhERD), as well as protein–ligand complex data including BindingMOAD (DiffSBDD-M) and CrossDock (DiffSBDD-C). Incorporation of domain knowledge was also heterogeneous, spanning molecular surface information (Delete, ShapeMol, ShEPhERD) and pharmacophoric constraints (ShEPhERD, PGMG). This representative set of baselines provides a comprehensive perspective for assessing how various factors influence performance across different scenarios. Detailed characteristics of all evaluated models are summarized in **Fig. 2a–b**.

**Fig. 2.**
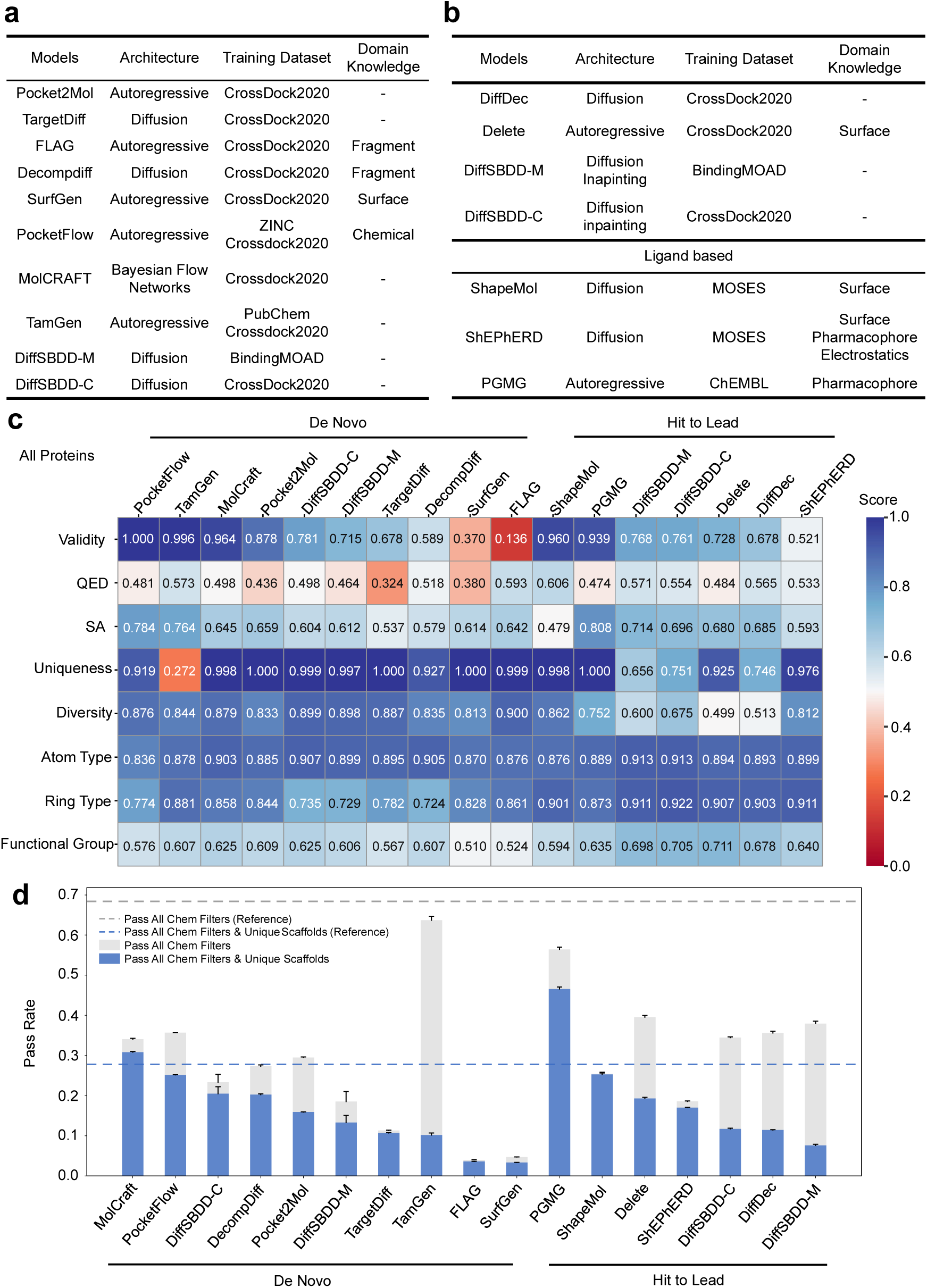
Overview of Evaluated Molecular Generative Models and Assessment of Core Performance Metrics. **a,** Summary of de novo molecular design methods and their key characteristics (architecture, training dataset, and integrated domain knowledge). **b,** Summary of H2L optimization methods and their key characteristics (architecture, training dataset, and integrated domain knowledge). **c,** Evaluation results of basic molecular properties, including validity, QED, SA score, uniqueness, diversity, and motif distributions(atom types, ring types, functional groups) with reference actives. Values represent the mean from three independent replicates; all metrics are normalized to a 0–1 scale, where higher values indicate better performance. **d,** Evaluation results of chemical safety (filter pass rates, reflecting compliance with industry-standard rules for avoiding unstable/reactive molecules) and Bemis-Murcko scaffold diversity among filter-passing generated molecules. Bars represent the mean ± standard deviation from three independent replicates.

In addition to comparing the distinctions among the models, we performed a fine-grained categorization of the test data. Since nearly all de novo SBMG methods included in our comparison were trained on CrossDock, except for DiffSBDD-M, we partitioned the collected data into two groups based on whether each protein appeared in CrossDock. These groups are designated as “Proteins in CrossDock” and “Proteins Not in CrossDock”, respectively, to analyze the impact of training data protein exposure on model performance. DiffSBDD-M was included primarily to evaluate whether training on crystal structure data significantly influences model behavior (vs. CrossDock’s simulated complexes). Unless otherwise specified, all statistical results presented in this study are based on three independent replicates.

### Assessment of Basic Generative Capabilities of Molecular Generative Models: From Standard Metrics to Chemical Safety and Scaffold Diversity

While the direct evaluation of generated molecules using common standard metrics does not directly reflect a method’s ability to produce bioactive compounds for a given target, it serves as an essential assessment of the structural and chemical plausibility of the outputs, which is a foundational prerequisite for any practical drug discovery application. As summarized in **Fig. 2c**, we compared all methods across five core standard performance metrics: validity (proportion of chemically intact molecules recognized by RDKit), QED (Quantitative Estimation of Drug-likeness, a composite score of drug-like properties), SA score (estimate of synthetic feasibility), uniqueness (proportion of non-duplicate molecules per target), and diversity (Tanimoto similarity-based coverage of chemical space). Additionally, following the approach of Lin et al.^42^, we quantified the distributional discrepancies between the generated molecules and reference active compounds across multiple structural hierarchies, including atom types, ring types, and functional groups, using Jensen–Shannon Divergence (JSD); scores were calculated as 1 – JSD, where 1 indicates perfect alignment with reference distributions. Detailed descriptions of all metrics are provided in the Methods section.

Among the de novo generation methods, a subset exhibited notably poor performance on validity. For instance, FLAG and SurfGen demonstrated substantially lower validity scores (< 0.4), indicating a failure to consistently produce chemically viable molecules and suggesting insufficient learning of fundamental molecular grammatical rules. In contrast, top-performing approaches such as PocketFlow and TamGen achieved validity scores approaching or equal to 1, underscoring the value of molecular pre-training in capturing essential syntactic constraints^1^. MolCraft, a BFN-based model, also delivered competitive validity (∼0.96), highlighting the utility of advanced architectural designs for ensuring chemical plausibility.

Subsequent analyses were restricted to valid molecules for further benchmarking. Within this subset, TamGen displayed markedly lower uniqueness (∼27% unique molecules), far below the >90% observed for other methods, indicating limited exploration of chemical space, likely due to over-reliance on pre-trained patterns of active molecules. Beyond uniqueness, all de novo models performed comparably on QED (∼0.4–0.6), SA (∼0.5–0.8), and diversity (∼0.8–0.9), demonstrating their ability to generate drug-like, structurally varied molecules. When aligned to reference actives, generated molecules showed closest alignment for atom types (score ∼0.8–0.9), followed by ring types (score ∼0.7–0.9), and the largest divergence for functional groups (score ∼0.5–0.6), a hierarchy that suggests models struggle to replicate the precise functional group patterns that mediate target binding.

For H2L approaches, the highest validity rates were consistently observed in models trained on large molecular datasets with less constraints: PGMG (ChEMBL-trained) and ShapeMol (MOSES-trained) achieved validity >75%, underscoring the critical role of extensive data in learning fundamental molecular grammar. A key distinction emerged between H2L subcategories: fragment-based methods (DiffSBDD, Delete, DiffDec) produced compounds with reduced structural diversity compared to de novo models and ligand-based H2L methods, due to the constraints imposed by the given initial fragment structure. In contrast, ligand-based H2L methods (ShEPhERD, PGMG, ShapeMol) generated more diverse molecules, as they lack such explicit fragment constraints and instead leverage features of reference actives to guide generation. Across most evaluated metrics, including SA, QED, and alignment of atom/ring/functional group distributions, H2L models generally outperformed de novo designs. Importantly, the “atom type > ring type > functional group” alignment hierarchy observed in de novo models were conserved in H2L approaches.

We further stratified performance by “Proteins in CrossDock” vs. “Proteins Not in CrossDock” (**SI-Fig. 2**). The observed trends are consistent with the statistical results obtained across the full set of targets, confirming results are not driven by target overlap with training data.

Beyond basic plausibility, practical drug discovery requires avoiding unstable, reactive, or promiscuous “high-risk” molecules; we therefore applied a panel of industry-standard chemical filters (such as Eli Lilly’s reactivity rules and Novartis Institute for BioMedical Research’s criteria and ChEMBL rules)^43,44^ to generated molecules and computed pass rates (**Fig. 2d**). Nearly 70% of reference active molecules passed all filters, whereas among de novo and H2L models, only TamGen and PGMG achieved pass rates exceeding 50%. Most other methods fell below 40%, with some (e.g., FLAG, SurfGen) below 5%. The relative advantage of TamGen and PGMG may be attributed to their training on large-scale datasets of known active molecules, which implicitly encode safety-related structural patterns. These results highlight that current generative models frequently produce high-risk compounds, which not only fail to advance drug discovery but also introduce unnecessary experimental risks, underscoring the need for rigorous post-generation filtering. We additionally analyzed Bemis-Murcko scaffold diversity among filter-passing molecules (**Fig. 2d**, Methods section). Among de novo models, TamGen exhibited high scaffold redundancy (>84% duplicate scaffolds), while two other autoregressive models (Pocket2Mol, PocketFlow) also showed substantial redundancy (>30% duplicates). In contrast, diffusion-based (e.g., TargetDiff, DiffSBDD) and BFN-based (MolCraft) models produced more diverse scaffolds; notably, MolCraft was the only de novo method where the proportion of unique valid scaffolds exceeded the reference set. For H2L models, fragment-based (fragment and pocket context constrained) methods yielded reduced scaffold diversity, while ligand-based methods (free of explicit fragment and pocket context constraints) achieved greater structural variety with PGMG being the only H2L method matching the reference set’s unique scaffold proportion. Stratified results for individual protein subsets are provided in **SI-Fig. 2**.

Collectively, these findings reveal a critical limitation: generating bioactive molecules requires not just correct constituent groups (atoms, rings, functional groups) but their proper compositional and spatial arrangement. However, current evaluation metrics fall short in effectively capturing a model’s capability in this regard. Therefore, addressing this gap by introducing novel, bioactivity-focused criteria motivate the focus of the subsequent section.

### Conformational Prediction Limitations of Current Molecular Generative Models: A Multi-Dimensional Evaluation

In the preceding analyses, we conducted a comprehensive evaluation of both categories of methods, assessing fundamental molecular properties. For most binding-pocket-based molecular generative models, there exists an implicit expectation beyond mere generation SMILES or 2D molecular graph: the simultaneous prediction of precise binding conformations for the generated molecules, critical yet underappreciated requirement for SBDD. However, current 3D molecular generative approaches often yield physically implausible molecular structures^35^. To quantitatively evaluate structural validity, we employed the PoseBusters package, a rigorously validated framework that performs a comprehensive suite of standard quality checks. PoseBusters evaluates both intramolecular plausibility (e.g., aromatic ring planarity, compliance with standard bond lengths/angles) and intermolecular compatibility (e.g., steric clashes between ligand and pocket residues). Detailed computational protocols are provided in the Methods section.

As illustrated in **Fig. 3a**, the results demonstrate a different performance across the evaluated methods. Among de novo approaches, MolCraft yields the highest validity score (0.964), with a corresponding PB-valid score of 0.783 after conformational quality checks, indicating that most of its generated valid molecules (∼81%) adopt structurally plausible conformations. In contrast, PocketFlow, which ranks second, achieves a perfect validity score of 1.000 but exhibits a markedly lower PB-valid score of 0.449, implying that fewer than half of its valid molecules have reasonable conformation. The remaining de novo methods not only show lower validity proportions (**Fig. 2c**) but also display substantially reduced PB-Valid score among valid molecules, with PB-Valid scores falling below half or even one-third of their validity values, exemplified by Pocket2Mol (0.878 vs. 0.388), SurfGen (0.370 vs. 0.103), and FLAG (0.137 vs. 0.001).

**Fig. 3.**
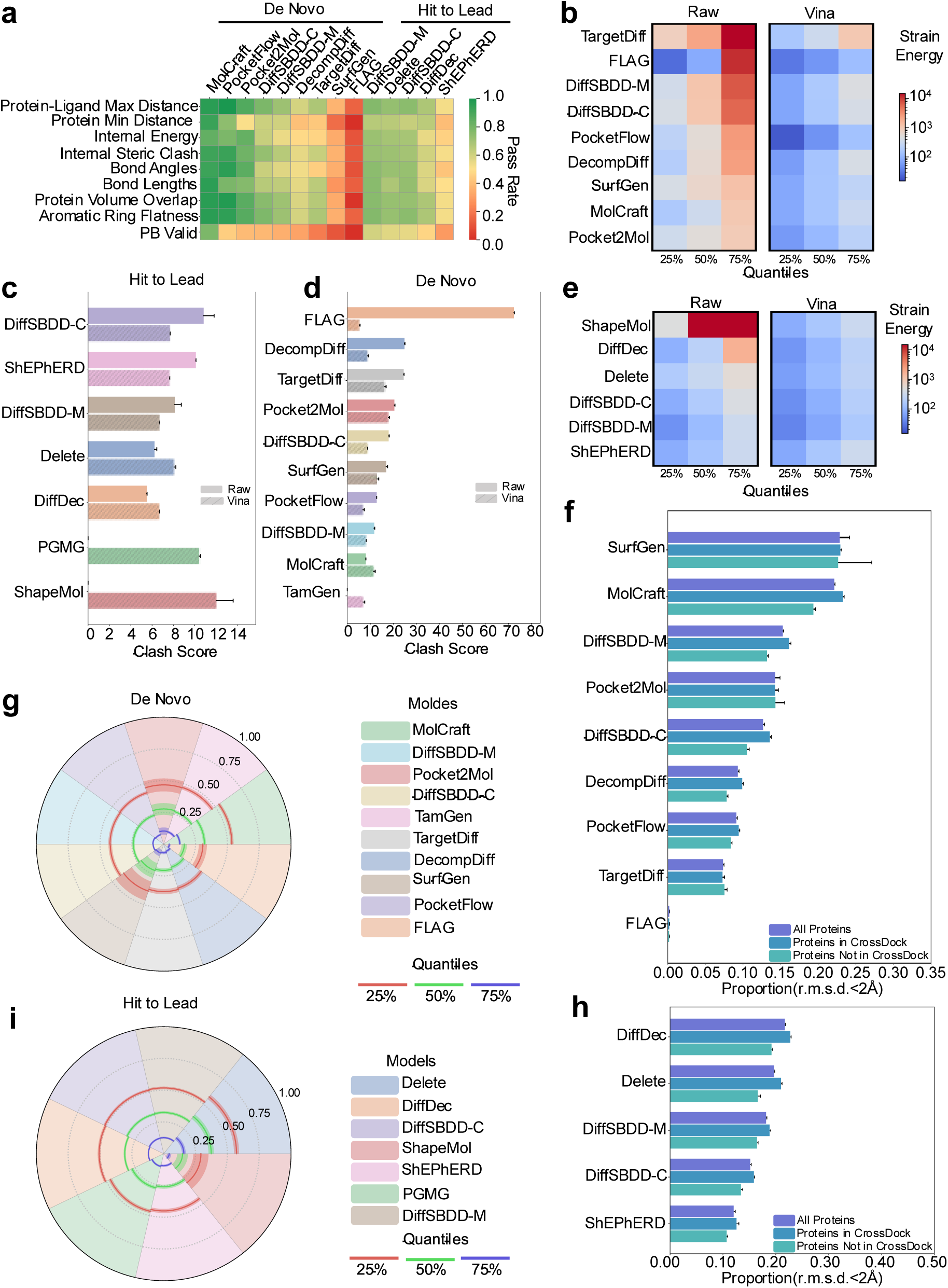
Multi-Dimensional Assessment of Conformational Quality in Structure-Based Molecular Generative Models. **a**, PoseBusters validation results for all generative models (de novo and H2L) across all protein targets. Molecules generated by ShapeMol were excluded from this evaluation due to the lack of guaranteed placement within the target binding pocket. **b**, Strain energy distribution of conformations generated by all de novo models across all protein targets. **c**, Steric clash frequency of conformations generated by all H2L optimization models across all protein targets. **d**, Steric clash frequency of conformations generated by all de novo models across all protein targets. **e**, Strain energy distribution of conformations generated by all H2L optimization models across all protein targets. **f**, Redocking root mean square deviation (r.m.s.d.) evaluation of all de novo models on proteins stratified by CrossDock training set inclusion (all vs. training-seen vs. training-unseen). **g**, Redocked interaction evaluation (vs. reference actives) of all de novo models across all protein targets. **h**, Redocking r.m.s.d. evaluation of all H2L optimization models on proteins stratified by CrossDock training set inclusion (all vs. training-seen vs. training-unseen). **i**, Redocked interaction evaluation (vs. reference actives) of all H2L optimization models across all protein targets. Notably, we calculated the PoseBusters pass rate using the total number of sampled molecules (not just chemically valid molecules) as the denominator. This design inherently classifies invalid or unsuccessfully sampled molecules as conformational failures, providing a holistic measure of performance from initial generation to conformational validation.

For H2L models (**Fig. 3a**), DiffSBDD-M achieved the highest overall pass rate (0.597), followed by Delete (0.563), DiffSBDD-C (0.552), and DiffDec (0.466). These models outperformed all de novo models except MolCraft, while the ligand-based method ShEPhERD (0.290) ranked last. These results demonstrate the limitations of relying solely on ligand features (vs. pocket and fragment context) for conformational prediction. Results stratified by targets in or not in CrossDock training set, provided in **Extended Fig. 1**, are consistent with the overall trend. Notably, nearly all methods exhibit lower PB-valid scores on proteins not in CrossDock training set compared to in ones, underscoring challenges in generalizing conformational prediction to novel targets.

Owing to the scarcity of experimentally determined protein–ligand complex structures, most current studies resort to using computationally generated docking poses as supervisory signals. Among various tools, AutoDock Vina (Vina) has become the de factor standard for constructing such datasets, including the widely adopted CrossDock set, due to its efficiency and accessibility. While this approach is both practical and necessary, it raises a fundamental question: does training a model within a conformational space “defined” by Vina’s scoring function and search algorithm inherently cap the model’s performance at the level of Vina’s own capabilities? To further investigate whether the use of such data limits a model’s ability to generate conformations that outperform those produced by Vina, we analyzed the strain energy (a measure of intramolecular stability) of binding conformations across different quantiles. As shown in **Fig. 3b**, for de novo methods, the directly generated conformations generally exhibit higher strain energies compared to those from Vina The relatively better performance of FLAG at the 25th quantile can be attributed mainly to two factors: its low number of valid molecules (reducing variability) and post-generation force field refinement. Although these results indicate that current de novo methods have not yet surpassed Vina in producing low-strain-energy conformations, several approaches, including MolCraft, DecompDiff and PocketFlow demonstrated competitive potential, suggesting that diverse architectural strategies hold promise for improved conformational modeling. For H2L models (**Fig. 3e**), the conformational strain energies of most approaches are closer to those produced by Vina compared to de novo design methods. This indicates that providing reference molecular information or initial fragments facilitates the generation of conformations with more realistic strain energy profiles.

Among H2L models, DiffSBDD-M (crystal-trained) outperformed DiffSBDD-C (simulation-trained), further emphasizing that increasing training data quality (experimentally derived vs. simulated) can improve conformational prediction accuracy.

Furthermore, in **Fig. 3c–d**, we quantified the average number of clashes between generated conformations and the protein pocket, a critical metric for binding feasibility, as severe steric overlaps directly preclude stable ligand–protein interactions and render such conformations biologically irrelevant. Notably, select advanced generative models surpassed Vina in minimizing these detrimental clashes: Among de novo models, MolCraft achieved the lowest clash frequency, outperforming Vina; for H2L models, Delete and DiffDec similarly exhibited lower clash frequencies than Vina.

This indicates that certain advanced deep learning methods can surpass the patterns inherent in the training data derived from Vina generated poses and delivering improved steric compatibility between ligands and proteins. Stratifying by CrossDock training set inclusion (**SI-Figs. 3–4**) showed no significant performance differences. The results align closely with those from the full dataset, suggesting that conformational strain or clash performance is less independent from prior target exposure.

Experimentally determining the binding conformations of all generated molecules is nearly infeasible. In the field of protein design, it has become a widely accepted computational validation practice, as exemplified by tools such as RFdiffusion^45^, to assess the rationality of generated conformations by comparing them against those predicted by specialized structure prediction tools like AlphaFold2^46^ and evaluating their consistency. For molecular generation, analyses based on strain energy and protein–ligand clashes indicate that traditional docking methods still outperform current generative approaches in conformational prediction accuracy. Therefore, using traditionally docked poses as reference structures and computing the r.m.s.d. between generated and reference poses can serve as a reasonable and practical proxy for evaluating the quality of generated conformations.

We evaluated both de novo and H2L models by quantifying the proportion of generated poses with r.m.s.d. < 2 Å relative to poses re-docked using Vina (detailed results in **Figs. 3f** and **3h**). Among the de novo methods, SurfGen performed best, with 22.9% of its poses achieved r.m.s.d. < 2 Å, followed by MolCraft (22.2%) and DiffSBDD-M (15.3%). In the H2L category, the top-performing method is DiffDec, with 21.8% of its poses showing r.m.s.d. < 2 Å, followed by Delete (19.8%). These results indicate that only a small fraction of the poses generated by current methods exhibit binding modes consistent with those derived from docking. Meanwhile, our analysis further examined the influence of protein presence in the training data. As shown in **Fig.3f** and **Fig.3h**, nearly all models exhibited weaker redocking performance on proteins unseen during training compared to those seen previously. This indicates that evaluating redocking performance solely on seen proteins may lead to an overestimation of model capability, underscoring the necessity of including unseen proteins in such assessments. However, the r.m.s.d. metric calculated in this manner is not an ideal indicator, as it is influenced by both the accuracy of the docking method and the flexibility of the molecules involved. In **SI-Fig. 5–6**, we analyzed the distributions of rotatable bonds and ring counts in molecules generated by all models in a single sampling round. Among de novo models, SurfGen produced molecules with the fewest rotatable bonds on average and the highest average number of rings (**SI-Fig. 5 and 6**), resulting in the lowest molecular flexibility. This structural rigidity reduced conformational complexity, facilitating more accurate pose prediction and thereby contributing to its improved redocking r.m.s.d. performance. A similar trend was observed for Pocket2Mol. Among H2L models, ShapeMol exhibited the highest average ring count and the fewest rotatable bonds. However, since its generated conformations are not guaranteed to be positioned within the binding pocket, ShapeMol was excluded from the redocking r.m.s.d. evaluation.

These findings further indicate that when using r.m.s.d. to evaluate a model’s ability to generate plausible conformations, caution is warranted against models potentially taking shortcuts by producing overly rigid molecular structures. Incorporating flexibility analysis (e.g., rotatable bond and ring count) into the evaluation of r.m.s.d. performance enables a more comprehensive and objective understanding of a model’s conformational prediction capability.

To complement conformational plausibility assessments, we further examined another critical requirement for active molecules: the ability to form key biological interactions (such as hydrogen bonds and π-stacking) comparable to those observed in reference active compounds by a new interaction evaluation framework. Previous methods such as PoseCheck commonly evaluate interaction similarity by computing the interaction fingerprint (IFP) similarity between generated poses and a single reference active’s docked structure. A modest improvement involves using multiple reference molecules and retaining the maximum IFP similarity between each generated molecule and any reference. However, these approaches suffer from two critical limitations: (1) docked poses inherently introduce inaccuracies due to intrinsic flaws in docking algorithm; and (2) reliance on a small number of references amplifies bias, as individual actives may not represent the full spectrum of functional interaction modes. To mitigate these issues, we designed a weighted, aggregate interaction scoring framework to reduce conformational uncertainty and enable more robust assessment of interaction fidelity.

First, for each target, we used Vina dock all reference active molecules to obtain their binding modes and used ProLIF^47^ to quantify the types and frequencies of interactions formed by these molecules. We then computed the occurrence rate of each interaction type across all reference compounds and retained only those types whose frequency exceeded a predefined threshold (10% in this work), to exclude rare, non-conserved interactions. Subsequently, we normalized the frequencies of the retained interaction types to derive an “importance score” for each type (sum of all scores = 1), with higher scores assigned to more frequently observed interactions (reflecting greater conservation and potential functional relevance). The final interaction score for a generated molecule was defined as the sum of the scores corresponding to each interaction it forms, resulting in a value ranging from 0 to 1 (details in Method section). This design offers two main advantages: (1) aggregating patterns across all reference docked poses and excluding low-frequency interactions mitigates errors from individual docking inaccuracies; and (2) weighting interactions by conservation avoids equating trivial and functionally critical interaction. To assess how well models replicate reference-like interaction patterns, we quantified the proportion of generated molecules whose interaction scores exceeded key quantiles (25th, 50th, 75th) of reference active scores.

All analyses used interaction scores computed after redocking generated molecules with Vina to ensure alignment with physiologically relevant binding modes. Results for de novo and H2L models are presented in **Figs. 3g** and **3i**, respectively. Among de novo models (**Fig. 3g**), MolCraft performed best: 53.5%, 32.0%, and 12.7% of its molecules exceeded the 25th, 50th, and 75th quantiles, respectively. These values were the highest in the de novo category but were substantially lower than the corresponding proportions for reference actives, highlighting a persistent gap in replicating functional interactions. Other competitive de novo models included TamGen, Pocket2Mol, PocketFlow, DiffSBDD-M and DiffSBDD-C. For H2L models (**Fig. 3i**), Delete delivered the strongest performance: 57.2%, 37.3%, and 15.8% of its molecules exceeded the 25th, 50th, and 75th quantiles, respectively. These values align more closely with the reference actives than those from the best de novo model, MolCraft. The next best performers were pocket-based methods such as DiffSBDD-M, DiffSBDD-C and DiffDec, while ligand-based approaches ranked lowest. This underscores the importance of explicitly incorporating pocket information for accurate interaction modeling. Among ligand-based methods, those integrating pharmacophore constraints (PGMG, ShEPhERD) outperformed ShapeMol (shape-only), confirming that pharmacophore awareness enhances capture of critical interaction patterns, even in the absence of pocket context.

To further assess the quality of interactions in the generated poses by these methods, we compared the interaction scores calculated between the generated poses and the docked reference poses, along with their statistical outcomes, and evaluated model performance across different protein subsets: proteins included in the CrossDock training set and those excluded (**SI-Fig. 7**). For both de novo and H2L models, whether using Vina docked poses or generated poses, nearly all methods that include protein information in input performed worse on proteins not in CrossDock than on proteins seen during training, reinforcing that performance on familiar targets overestimates real-world utility. Furthermore, when comparing pose sources, different methods exhibited distinct behaviors. Among de novo models, MolCraft, Pocket2Mol, TargetDiff, DiffSBDD-M, and DiffSBDD-C achieved higher interaction scores with their generated poses than with Vina docked poses across both protein subsets. Similarly, H2L models, Delete and DiffDec showed the same trend. In contrast, the remaining methods attained higher interaction scores using Vina docked poses compared to their original generated poses, indicating their generated conformations lack intrinsic interaction fidelity. These observations suggest that, at the level of interaction modeling, certain methods may have surpassed the capability of classical tools such as Vina, representing a promising step toward more accurate SBMG.

### Limited Capacity of Current Molecular Generative Models to Rediscover Known Active Compounds: A Target-Stratified Evaluation

By moving beyond error-prone computational proxies like docking scores, we directly assessed the models’ ability to rediscover experimentally validated active compounds and scaffolds across diverse protein targets. This evaluation directly reflects a model’s utility in hit discovery, as ligand identification for a given target relies on two complementary computational paradigms: virtual screening (ranking existing library molecules by predicted activity) and de novo design (generating novel structures via learned ligand–target interaction rules). While their methodologies differ, both aim to identify potent ligands that bind the target of interest. Virtual screening offers a practical advantage: hit compounds identified from commercial libraries can be rapidly acquired for experimental validation, facilitating the direct assessment of the screening protocol. In contrast, molecules generated through de novo design typically require de novo synthesis, which presents a significant barrier to experimental confirmation. Owing to this synthetic bottleneck, the field often relies on computational docking scores (e.g., Vina scores) as a proxy metric to prioritize generated molecules with predicted high activity. However, the well-documented high false-positive rate associated with these scoring functions severely limits their reliability in gauging a model’s true potential for identifying genuine binders^48–50^. To overcome this limitation and provide a more rigorous evaluation, we curated an extensive set of known active compounds across diverse protein targets. This benchmark enables the direct assessment of a generative model’s capacity to discover active ligands, offering a realistic measure of its utility in hit discovery stage. We first evaluated de novo generative models for their ability to recover known active molecules (hit recovery) and Bemis-Murcko scaffolds (scaffold recovery) across the 120 targets. To dissect model performance on familiar versus novel targets, which is a key measure of real-world utility, we further stratified performance by “Proteins in CrossDock” vs. “Proteins Not in CrossDock”

As shown in **Fig. 4a** and **4c**, while most methods struggled to generate exact active molecules in both settings, the majority succeeded in reproducing Bemis-Murcko scaffold across many targets. This suggests that current approaches capture only coarse-grained patterns of protein-ligand interactions, failing to model the precise steric and electronic complementarity required to generate specific R-group decorations.

**Fig. 4.**
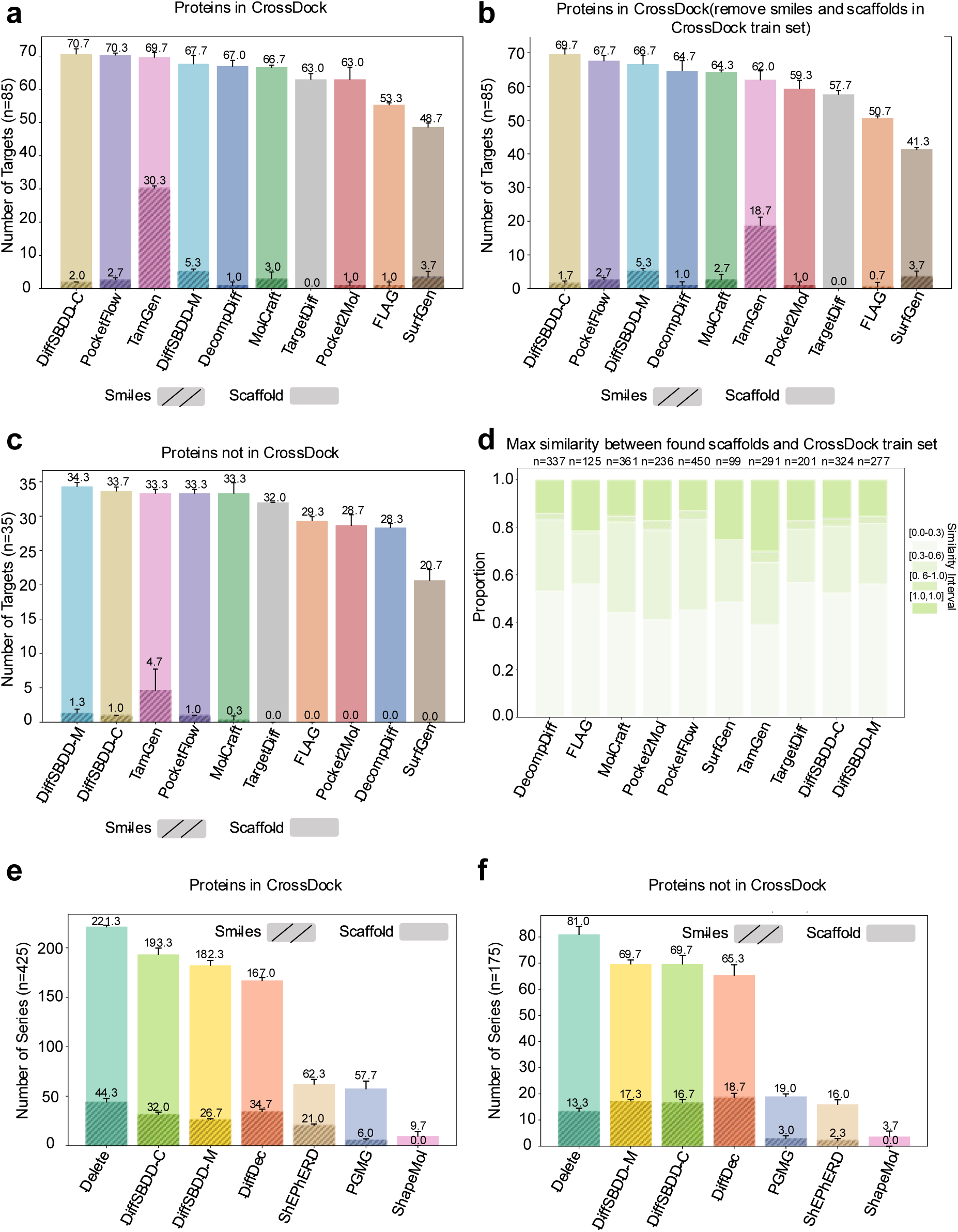
Evaluation of Molecular Generative Models for Identifying Active Molecules and Scaffolds. **a**, De novo models on proteins in CrossDock (n=85). **b**, De novo models on proteins in CrossDock (n=85), after excluding generated molecules/scaffolds overlapping with the CrossDock training set. **c**, De novo models on proteins not in CrossDock (n=35). **d**, Maximum similarity between active scaffolds from de novo models (proteins in CrossDock, n=85) and CrossDock training set scaffolds. **e**, H2L models on compound series for proteins in CrossDock (n=425). **f**, H2L models on compound series for proteins not in CrossDock (n=175). Note all evaluations assess model ability to recover active molecules and active Bemis-Murcko scaffolds, where histogram bars represent the mean number of targets where models recovered at least one active molecule (hatched bars) or at least one active Bemis-Murcko scaffold (solid bars), across three independent replicates. Error bars denote the standard deviation of the mean, reflecting variability in performance across replicates. Data are stratified by whether protein targets were included in the CrossDock training corpus; “n” denotes the number of protein targets (for de novo models) or compound series (for H2L models) in each subset. Notably, the H2L model DiffDec requires predefined atomic anchors to identify modification sites. These anchors are derived by aligning the common substructure of the compound series to reference actives; shared modification sites across reference actives are weighted by their occurrence frequency, and molecule generation is guided by these sites.

On Proteins in CrossDock (**Fig. 4a**), only TamGen recovered exact active molecules for more than a handful of targets; all other methods showed minimal success (recovering actives for 0–6 targets). To test if performance stemmed from memory of training data (rather than learned generalizable rules), we re-evaluated hit recovery after removing any generated molecules/scaffolds overlapping with CrossDock’s training set (**Fig. 4b**). Most models exhibited performance declines: TamGen’s active molecule recovery dropped from 30.3 to 18.7 targets, while scaffold recovery rates decreased across all methods (with DiffSBDD-C and PocketFlow remaining top performers). FLAG and SurfGen showed particularly poor scaffold recovery post-deduplication. For exact molecule recovery, only DiffSBDD-M, SurfGen, MolCraft, and PocketFlow achieved modest success (recovering actives for 3–5 targets).

We further computed the maximum Tanimoto similarity between generated scaffolds and scaffolds in CrossDock training data (**Fig. 4d**). Autoregressive models, particularly TamGen, SurfGen, FLAG, PocketFlow and Pocket2Mol produced numerous scaffolds same as training examples and analogs highly similar to training examples, with TamGen generating many nearly identical or identical replicates of known molecules (**SI-Fig.8**), confirming that their performance on proteins in CrossDock is inflated by memory effects. On Proteins Not in CrossDock (**Fig. 4c**), half of the de novo models failed to recover any active molecules, a stark contrast to the Proteins in CrossDock subset, where only TargetDiff showed complete failure. This highlights substantial overfitting in current de novo models and limited generalization to novel targets. Under this configuration, TamGen still recovered active molecules for the highest number of protein targets, demonstrating the unique advantage of its pre-training on a large corpus of active molecules.

To complement our de novo model analysis, we evaluated the capability of H2L methods to recover active molecules/scaffolds. Compared to de novo design, the provision of molecular fragments effectively constrains the search space and offers critical structural context to guide bioactive generation. As illustrated in **Fig. 4e–f**, fragment-based H2L models (incorporating protein pocket and molecular fragment information) outperformed ligand-based methods (relying solely on active ligand features) across both target subsets. Among fragment-based approaches, the autoregressive model Delete achieved the highest performance, successfully recovering active molecules in 44.3 series and scaffolds in 221.3 series corresponding to proteins in CrossDock. Within the diffusion-based frameworks, DiffDec exhibited superior results, retrieving active molecules in 34.7 series and scaffolds in 167.0 series associated with known targets. DiffSBDD, which employs a molecular inpainting strategy, demonstrated considerable optimization capability, highlighting both the advantage of inpainting-based approaches and the broader potential of integrating such techniques with de novo design for molecular optimization. Notably, DiffSBDD-C, trained on a larger set of diverse small molecules, demonstrates a discernible advantage over DiffSBDD-M which was trained exclusively on crystal structures when evaluated on proteins in CrossDock. This result underscores the effectiveness of expanding molecular diversity in enhancing molecular optimization performance. Among ligand-based methods, approaches incorporating pharmacophore constraints (ShEPhERD, PGMG) outperformed those relying exclusively on shape information (ShapeMol). This is likely attributable to the richer chemical information and implicit receptor-specific features captured by pharmacophore representations. Furthermore, ShEPhERD significantly surpassed PGMG, potentially due to its integration of multimodal information and effective leveraging of such signals to guide molecule generation.

On Proteins Not in CrossDock, trends mirrored those in the training-seen subset. However, diffusion-based H2L models (DiffDec, DiffSBDD) exceeded autoregressive Delete in active molecule recovery. This highlights the superior generalization of diffusion frameworks for optimizing molecules against novel targets, a critical advantage for real-world drug discovery where many targets lack extensive training data.

In evaluating structure-based molecular generative models, we should note that the “active compound rediscovery task” has inherent limitations that necessitate cautious interpretation of results. First, the chemical space is vastly larger than current experimental capabilities can explore; the 220,005 experimentally confirmed active molecules spanning 120 protein targets curated in this study represent a biased subset constrained by specific technologies and data sources, failing to capture the full distribution of a target’s bioactive space^51,52^. Second, model generation is constrained by architectural biases and sampling scales, precluding unbiased navigation of bioactive space. Thus, a model’s failure to recapitulate known actives for a specific target may stem from sampling biases in the known active set or inherent exploration limitations of the model, rather than indicating inactive generated molecules or poor method performance. Notably, this study mitigates such uncertainties through multi-target statistics, stratified evaluation (proteins in vs. out of CrossDock), and multi-metric integration. If a method consistently fails to rediscover active molecules across most targets and independent replicates, it can still reflect a systematic limitation in its ability to navigate bioactive chemical space.

### Limited Screening Efficiency of Current Molecular Generative Models: A Hit Rate-Based Evaluation

In the preceding section, we examined whether these generative methods could successfully rediscover active molecules for their respective targets. However, this rediscovery-focused evaluation does not capture a critical practical attribute: the efficiency with which models identify active compounds in real-world applications. In virtual screening, the hit rate is a gold-standard metric, serves as a robust indicator of the efficiency with which active compounds are identified within a single screening campaign^53^. This metric is of critical importance in real-world hit discovery efforts.

To systematically characterize the efficiency of each model in discovering active entities, we quantified the hit rate of each model for both active molecule and active scaffold discovery. The hit rate is defined as the ratio of unique active entities (molecules or scaffolds) recovered to the total number of molecules sampled, as formalized in Equation (1):

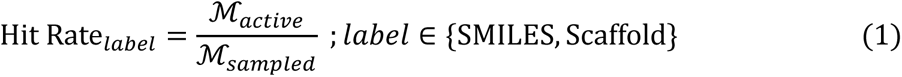

Here, ℳ*_active_* denotes the number of unique active molecules or molecules bearing active Bemis-Murcko scaffolds identified in the sampled set, while ℳ*_ref_* represents the total number of generated molecules sampled for evaluation. Compared to the use of computational metrics for evaluating the quality of generated molecules, directly utilizing a collection of known active molecules as a reference is not susceptible to biases inherent to the computational methods themselves.

As shown in **Fig. 5a-b**, de novo generative models exhibited remarkably low hit rates for active molecule discovery. On average, the highest molecular hit rate among these models was only 0.124% when evaluated on proteins included in the CrossDock training set, and this value dropped to just 0.024% for proteins not in CrossDock. In contrast, the hit rate for active scaffold discovery was more than 10-fold higher across all de novo models, highlighting that generating pharmacologically plausible scaffolds is substantially less challenging than generating fully active molecules with precise steric and electronic complementarity to the target pocket. Nevertheless, progressing from identified active scaffolds to functionally active molecules remain a major bottleneck for current de novo methodologies.

**Fig. 5.**
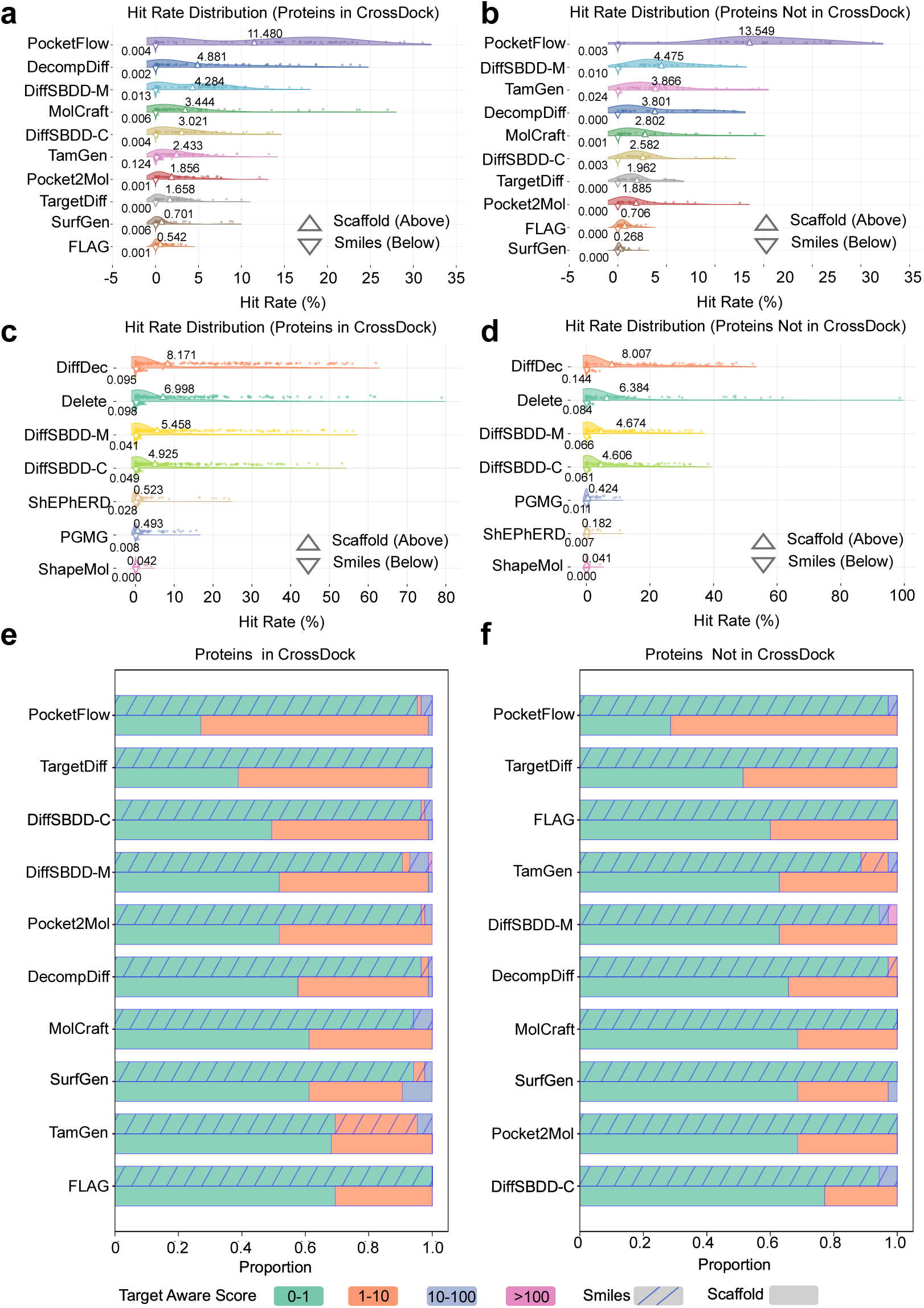
Comprehensive Evaluation of Screening Efficiency and Target-Awareness in Molecular Generative Models. **a**, Hit rate distribution of de novo generative models for proteins included in the CrossDock training set. **b**, Hit rate distribution of de novo generative models for proteins not included in the CrossDock training set. **c**, Hit rate distribution of H2L optimization models for proteins included in the CrossDock training set. **d**, Hit rate distribution of H2L optimization models for proteins not included in the CrossDock training set. **e**, Evaluation of target-aware capabilities in de novo generative models for proteins included in the CrossDock training set. **f**, Evaluation of target-aware capabilities of de novo generative models for proteins not included in the CrossDock training set. Notably, the H2L model DiffDec requires predefined atomic anchors to identify modification sites. These anchors are derived by aligning the common substructure of the compound series to reference actives; shared modification sites across reference actives are weighted by their occurrence frequency, and molecule generation is guided by these sites.

A direct comparison between **Fig. 5a** and **5b** further reveals that all de novo models exhibited reduced efficiency in active molecule discovery when tested on proteins outside the CrossDock training set; most models also showed declines in active scaffold hit rates for these unseen targets. Notably, among de novo models, PocketFlow demonstrated the most consistent performance across both seen and unseen targets, with active scaffold hit rates exceeding 10% overall and reaching over 30% for certain targets. This robustness underscores the value of its integrated molecular pre-training (on the ZINC database) in enhancing generalization to novel targets. For TamGen, while its scaffold-level hit rate was notably low, it achieved the highest molecular hit rate among de novo models in both test scenarios. However, its molecular hit rate was significantly lower on proteins not in CrossDock compared to those present in CrossDock, further suggesting potential data leakage during training.

To dissect the impact of model architecture on screening efficiency, we focused on de novo models trained exclusively on the CrossDock dataset (DecompDiff, MolCraft, DiffSBDD-C, Pocket2Mol, TargetDiff, SurfGen, and FLAG). Across both seen and unseen proteins, the top three performers in terms of hit rate were consistently either diffusion-based models or those leveraging BFNs. In contrast, autoregressive models (e.g., Pocket2Mol, SurfGen, FLAG) clearly underperformed on this metric. These results suggest that, for de novo molecule generation, further integration of molecular pre-training with advanced diffusion modeling holds promise for enhancing the efficiency of discovering active molecules and scaffolds.

We extended this hit rate evaluation to the H2L optimization task. As shown in **Fig. 5c–d**, fragment-based H2L models (which incorporate both fragment constraints and protein pocket context) exhibited a distinct advantage in hit rate relative to ligand-based counterparts (which rely solely on structural features of known active ligands). This finding underscores the superiority of directly integrating target structural context and fragment information over relying exclusively on ligand-derived features for optimization. Furthermore, among models specifically designed for H2L optimization, the diffusion-based DiffDec outperformed the autoregressive Delete; inpainting-based methods (e.g., DiffSBDD-C, DiffSBDD-M) also demonstrated strong performance.

Compared to de novo models, fragment-based H2L models (DiffDec, Delete, DiffSBDD-C, DiffSBDD-M) achieved significantly higher hit rates for active molecules. However, their scaffold-level hit rates were lower than those of the best de novo model (PocketFlow). This trade-off arises because H2L models are tasked with growing or modifying a predefined fragment, confining chemical space exploration to regions adjacent to the fixed scaffold and thereby reducing the diversity of generated scaffolds relative to de novo design.

### Limited Coverage of Bioactive Chemical Space by Molecular Generative Models: Hit Fraction Evaluation and Inference-Time Sampling Analysis

While the hit rate metric effectively quantifies the efficiency of identifying active compounds or scaffolds, it fails to capture two critical dimensions of practical relevance: the structural diversity of the discovered active entities and their coverage relative to the full known bioactive chemical space of a target. Evaluating a model’s ability to explore this bioactive space comprehensively is essential for assessing its utility in real-world drug discovery, where the goal is to identify a broad spectrum of viable candidates, not just a small subset. To address this gap, we propose the hit fraction metric to quantify the extent to which a model probes the active chemical space of a target. The hit fraction is defined as the ratio of unique active molecules or scaffolds recovered by a model to the total number of reference active molecules or scaffolds for that target, as formalized in Equation (2):

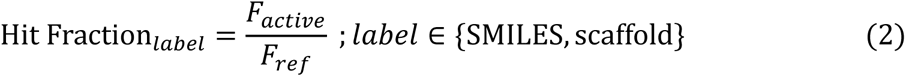

Here, 𝐹*_active_* denotes the number of unique active molecules or Bemis-Murcko scaffolds identified in the model’s sampled outputs, while 𝐹*_ref_* represents the total number of unique reference active molecules or scaffolds for the specific target protein. This metric directly reflects a model’s ability to “cover” the known bioactive space, complementing the hit rate’s focus on efficiency.

We first applied the hit fraction metric to assess de novo generative models across targets stratified by their presence in the CrossDock training set. As shown in **Extended Fig. 2a–b**, de novo models exhibited exceedingly low coverage of the known active chemical space for both molecules and scaffolds. For targets included in CrossDock, TamGen achieved the highest mean hit fraction for molecules (0.067%), while PocketFlow led for scaffolds (1.008%). To disentangle true generalization from training data memorization, we re-evaluated performance after removing any generated molecules or scaffolds that overlapped with the CrossDock training set (**Extended Fig. 2d**). This deduplication caused a notable performance drop: the maximum molecular hit fraction fell to 0.021% (TamGen), and the scaffold hit fraction decreased to 0.849% (PocketFlow). Critically, the post-deduplication hit fraction of de novo models was substantially lower than their performance on targets *not* in CrossDock (**Extended Fig. 2b** vs. **2d**). A parallel trend was observed for the hit rate metric (**Extended Fig. 2c** vs. **Fig. 5a**). These findings suggest that the models may have tended to memorize seen active samples during training, leading to an overestimation of performance while discouraging exploration of a broader bioactive space. This result underscores the need to develop improved model architectures and training strategies to overcome this limitation.

As shown in **Extended Fig. 2e-f,** we conducted a corresponding evaluation of H2L models. In this test scenario, Delete and DiffDec delivered the strongest performance. Delete consistently identified a greater number of distinct active scaffolds across different protein subsets, while DiffDec excelled at discovering active molecules on novel proteins. These were followed by the inpainting-based methods DiffSBDD-C and DiffSBDD-M. Among ligand-based methods, ShEPhERD achieved the best overall performance, followed by PGMG, with ShapeMol yielding the least favorable results. This trend underscores the practical value of incorporating richer prior information to enhance model efficacy in lead optimization scenarios.

Several noteworthy observations emerged: the proportion of active molecules and scaffolds identified by H2L models relative to the entire known active chemical space was substantially higher than that of de novo generation models. Furthermore, in contrast to the results from de novo design models, the hit fraction values for these optimization approaches exceeded their hit rate values. This indicates that H2L methods can more effectively explore the target-active chemical space by leveraging the information from the given fragment. Moreover, we observed that fragment-based H2L models achieved higher hit fraction values on proteins not in CrossDock than on those present in CrossDock. This result indicates that, when provided with an initial fragment, these models can leverage both protein pocket information and fragment constraints to achieve enhanced generalization capability.

Owing to computational constraints, our initial baseline evaluations sampled 1,000 molecules per target for de novo models and 200 molecules per compound series for H2L models. Recent studies, particularly in the domain of deep learning and large language models, have highlighted the presence of inference-time scaling laws^54^ where allocating greater computational resources during inference can lead to improved model performance to test whether this phenomenon applies to molecular generative models, we selected computationally efficient models from each category (de novo: TamGen, PocketFlow, MolCraft, DiffSBDD-C; H2L: DiffDec, DiffSBDD-C, ShapeMol, PGMG) and evaluated them across three distinct target systems. We sampled up to 100,000 molecules per target/series to assess whether increased sampling enhances the number of active scaffolds/molecules discovered and their hit rates (detailed procedures in Methods).

As shown in **Extended Fig. 3a**, all four de novo generative models produced a greater number of active scaffolds with increased sampling. However, TamGen consistently underperformed relative to the other three methods across all targets. The efficiency of bioactive scaffold discovery diminishes with increased sampling for all models, revealing a fundamental exploration bottleneck. This bottleneck is most severe for TamGen, suggesting its generative landscape is less conducive to navigating diverse regions of bioactive chemical space. **Extended Fig. 3b** demonstrates that all evaluated de novo generative model exception of TamGen and PocketFlow, which failed to yield any active molecules in one target, showed an increase in the number of active molecules with larger sample sizes. A comparison between **Extended Fig. 3a** and **b** reveals orders-of-magnitude differences between the number of active scaffolds and molecules identified at comparable sampling levels, underscoring the substantially greater difficulty of molecule-level discovery.

Similar trends were observed for H2L models (**Extended Fig. 3c–d**): increased sampling improved the absolute number of active scaffolds and molecules recovered, but with diminishing efficiency. When comparing across tasks, MolCraft (BFN-based) maintained competitive performance in de novo evaluations, demonstrating the potential of BFN architectures for exploring bioactive space. DiffSBDD-C (inpainting-based) performed consistently well in H2L tasks, highlighting the utility of inpainting strategies for constrained optimization. Critically, while increasing sampling volume elevated the absolute number of active entities recovered, it did not improve their enrichment rate, evidenced by the declining slopes of the curves in **Extended Fig. 3**. Furthermore, current generative models and scoring functions remain limited in their ability to prioritize active molecules reliably. Thus, indiscriminately scaling up sampling volume offers little practical value for real-world drug discovery, as it fails to address the core limitation: the inability to distinguish active from inactive molecules efficiently. This finding underscores the urgency of developing more accurate protein–ligand interaction prediction tools to guide candidate prioritization.

### Insufficient Utilization of Protein Structural Information in De novo Generative Models: Evaluation via the Target-Aware Score (TAScore)

While the preceding section assessed de novo molecular generative models’ ability to recover active molecules and scaffolds, a critical gap remains: activity alone does not guarantee binding selectivity, which is a key requirement for safe, clinically viable drugs. Ideally, SBMG should leverage unique features of a target protein’s binding pocket (e.g., size, residue composition, electrostatic potential) to produce molecules with target-specific binding, rather than generic structures that may interact with off-target proteins. Conventional molecular property metrics, along with the active recovery rate and scaffold recovery rate discussed in the previous section, fail to capture a method’s ability to perceive and utilize target-specific information. Therefore, in this section, we propose a new evaluation metric, the Target-Aware Score (TAScore) to assess the target-aware capability of generative methods. The TAScore measures the model’s ability to perceive target-specific information by comparing the ratio of molecule/scaffold recovery on a specific target to the background recovery ratio across all targets. The formal definition formalized in Equation (3):

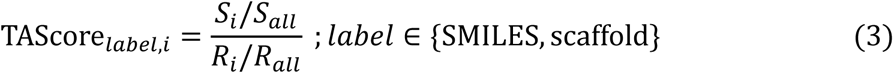

For the *i*-th target: 𝑅*_all_* is the total number of distinct molecules generated by the model across all 120 targets. 𝑅*_i_* represents the subset of 𝑅*_all_* that matches known active molecules or active scaffolds for target *i*. A large 𝑅*_i_* indicates that the model tends to generate active molecules or scaffolds for target *i* even when not conditioned on target i, signaling poor target specificity. 𝑆*_all_* is the total number of distinct molecules generated by the model when explicitly conditioned on target *i*. 𝑆*_i_* is the subset of 𝑆*_all_* that matches known active molecules or active scaffolds for target *i*. Higher TAScore values indicate stronger target awareness.

We applied the TAScore to assess target awareness across all de novo models, stratifying targets by their presence in the CrossDock training set **Fig. 5e–f**). At the scaffold level (TAScore_scaffold_), PocketFlow exhibited the strongest target sensitivity: For proteins included in the CrossDock training set, only 23 targets (27.06%) had a TAScore_scaffold_ < 1 (indicating no target awareness). In contrast, 38.82% of targets for TargetDiff (the second-highest performer) and 49.41% for DiffSBDD-C (third) had TAScore_scaffold_ < 1. All other de novo models failed to demonstrate target awareness (TAScore_scaffold_) for >50% of CrossDock-included targets. For proteins *not* included in the CrossDock training set, PocketFlow maintained robust performance: only 28.57% of targets had TAScore_scaffold_ < 1. All other models showed TAScore_scaffold_ < 1 for >50% of novel targets, highlighting PocketFlow’s unique ability to perceive and utilize target-specific information

At the molecular level (TAScore_smiles_), target awareness was substantially weaker across all models, reflecting the greater challenge of generating fully active, target-specific molecules (vs. scaffolds): For CrossDock-included proteins: TamGen achieved TAScore_smiles_ > 1 for 26 targets (30.59%), the highest among all models. DiffSBDD-M, MolCraft, and SurfGen followed, with TAScore_smiles_ > 1 for 8, 5, and 5 targets (9.41%, 5.88%, and 5.88%), respectively. All other models had TAScore_smiles_ > 1 for <5 targets. For novel proteins (not in CrossDock): Only six models demonstrated minimal target awareness: TamGen (4 targets), DiffSBDD-M (2), DiffSBDD-C (2), MolCraft (1), SurfGen (1), and PocketFlow (1). Notably, previous analyses suggest that the moderately better performance on proteins in CrossDock, particularly by methods such as TamGen may be attributed to their tendency to produce known active molecules or close analogues repeatedly. This is consistent with its sharp performance drop on novel targets.

Collectively, these results reveal a fundamental limitation of current de novo generative models: despite their design intent to leverage protein structural information, most fail to generate target-specific molecules. Instead, they often generate identical or highly similar scaffolds and molecules across different targets, a behavior that is undesirable in practical applications, as it may lead to substantial off-target effects. Given that H2Lmodels are constrained to generate compounds based on given fragment of known actives, this analysis was not applied to them.

### Evaluating H2L Optimization Models: Introducing the Mean Normalized Affinity (MNA) Score and Assessing Capability vs. Generalizability

A central contribution of this work is the establishment of a dedicated benchmark dataset for molecular optimization. While preceding analyses in this study assessed the hit rate and hit fraction performance of both de novo generative and H2L models, the latter are often explicitly intended to identify molecules exhibiting better bioactivity. To assess this capability, we introduce the Mean Normalized Affinity (MNA) Score to quantify the ability of H2L models to discover compounds with superior potency.

The MNA Score calculation proceeds in two steps, first normalizing experimentally validated binding affinities within each compound series and then computing a mean normalized value across all generated active molecules. Formally:

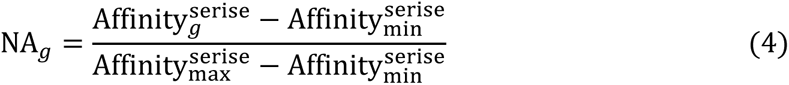

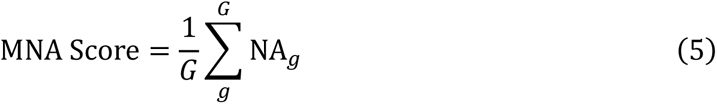

In Equation (4), 𝑔 denotes a generated active molecule for a specific compound series. 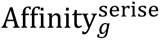 represents the binding affinity of molecule 𝑔 for its target. 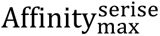 and 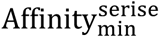 are the maximum and minimum binding affinities of reference active compounds within the same series, respectively. This normalization scales affinity values to a unified range [0, 1] across all series, enabling direct cross-series comparison. In Equation (5): 𝐺 is the total number of unique active molecules identified by a model across all compound series. NA_𝑔_ is the normalized affinity of molecule 𝑔. The MNA Score thus provides an aggregated measure of how well a model’s generated actives perform relative to the full range of reference potency for each series, with higher values indicating a greater proportion of potent generated molecules.

To assess H2L models’ optimization capability, we compared their performance across three independent replicates, excluding ShapeMol, which failed to generate any active molecules and thus could not be evaluated.

As shown in **Fig. 6a**, DiffDec emerged as the top performer: it achieved the highest total number of active hits (121.7) and the highest MNA Score (0.523), demonstrating its ability to generate both a greater quantity of active molecules and compounds with superior bioactivity. Delete ranked second, with 104.7 active hits and an MNA Score of 0.482. Next were the diffusion-based inpainting methods DiffSBDD-C (60.3 hits, MNA Score = 0.404) and DiffSBDD-M (56.3 hits, MNA Score = 0.415), followed by the ligand-based models ShEPhERD (26.0 hits, MNA Score = 0.465) and PGMG (11.0 hits, MNA Score = 0.453).

**Fig. 6.**
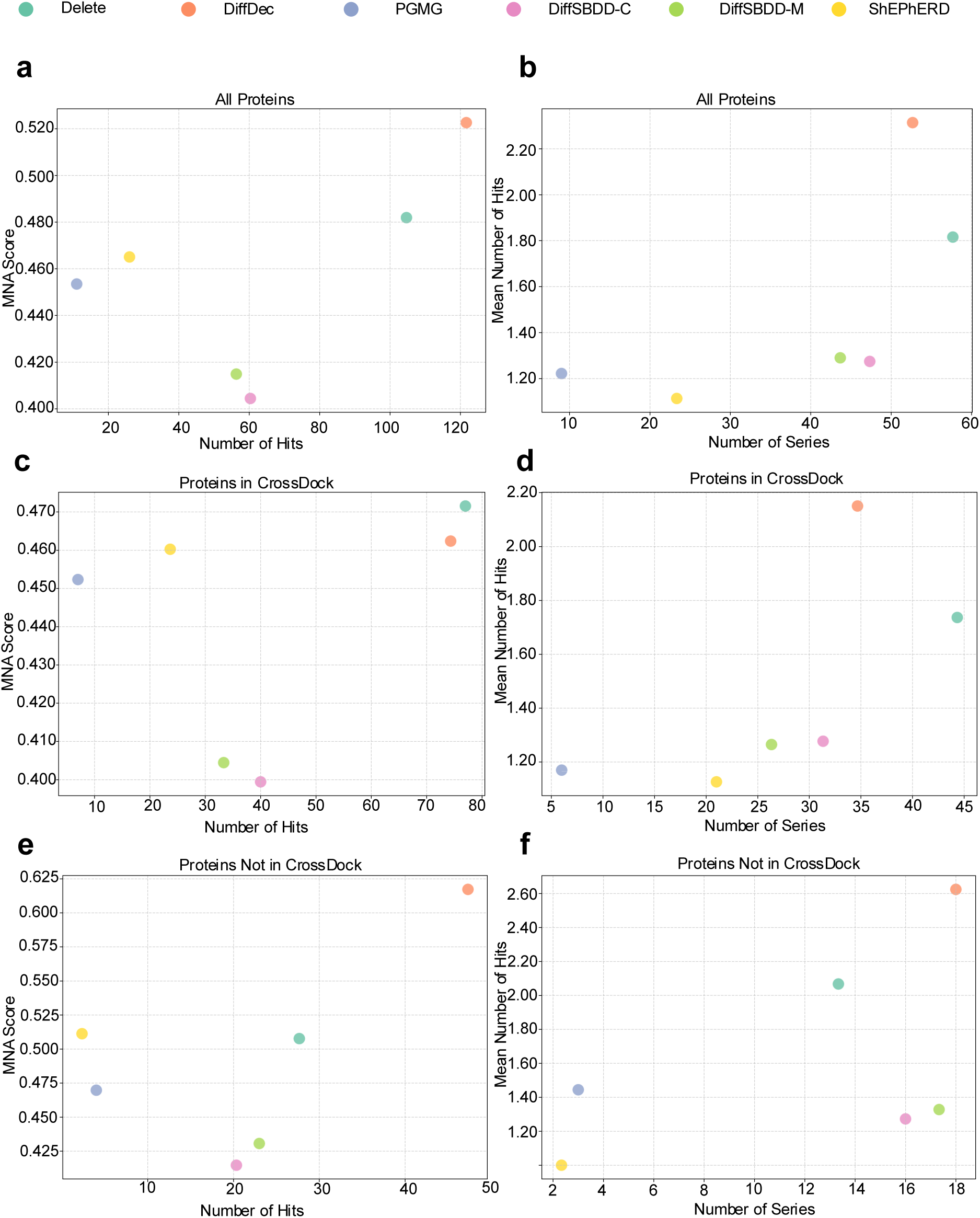
Evaluation of Hit-to-Lead (H2L) Optimization Models: Active Molecule Generation Potency and the Impact of Training Set Overlap with Target Proteins. **a, c, e**: Number of active compound hits recovered by H2L optimization methods, paired with their corresponding Mean Normalized Affinity (MNA) Scores. Evaluated across all protein targets (**a**), those included in the CrossDock training set (**c**), and those not included in the CrossDock training set (**e**). **b, d, f:** Number of compound series with at least one active hit recovered by H2L optimization methods, alongside the average number of active hits identified per series. Evaluated across all protein targets (**b**), those included in the CrossDock training set (**d**), and those not included in the CrossDock training set (**f**). Notably, the H2L model DiffDec requires predefined atomic anchors to identify modification sites. These anchors are derived by aligning the common substructure of the compound series to reference actives; shared modification sites across reference actives are weighted by their occurrence frequency, and molecule generation is guided by these sites.

Notably, ShEPhERD and PGMG exhibited higher MNA Scores than DiffSBDD-C/M despite fewer total hits. This suggests that while ligand-based models (which lack explicit protein pocket or fragment constraints) have lower odds of identifying active compounds, their integration of pharmacophore-based priors enables them to prioritize more potent molecules when actives are recovered.

Subsequently, we conducted a more detailed analysis of the number of series with hits and the average number of hits per series across different methods. As shown in **Fig. 6b**, Delete identified active compounds in 57.67 series with an average of 1.82 hits per series, followed by DiffDec, which detected actives in 52.67 series with an average of 2.31 hits. All other models performed lower across these metrics. Overall, models specifically tailored for fragment-based lead optimization scenarios demonstrated clear advantages in these aspects. Diffusion-based inpainting approaches, DiffSBDD-M and DiffSBDD-C also delivered competitive results. Notably, these fragment-based methods identified active molecules across a greater number of series compared to ligand-based approaches, further underscoring the importance of incorporating pocket and fragment information. Moreover, a comparison between DiffSBDD-M and DiffSBDD-C revealed that the former, trained on crystal structures, generated molecules with slightly higher activity, whereas the latter, trained on simulated data, succeeded in identifying actives across more series. Among ligand-based methods, ShEPhERD, which leverages more comprehensive prior information, achieved the best performance by identifying active molecules in more series and with higher potency, followed by PGMG, which incorporates pharmacophore constraints with training on ChEMBL dataset. In contrast, ShapeMol, which relies solely on molecular shape, failed to yield any active molecules. These findings suggest that different types of data and prior knowledge may enhance distinct capabilities of the models. Integrating simulated data, crystallographic data, and prior information holds promise for further improving model performance in lead optimization scenarios.

Finally, we further examined the influence of protein presence in the training set on model performance, as detailed in **Fig. 6c–f**. As shown in **Fig. 6c–d**, results on proteins in CrossDock were largely consistent with those across all proteins. On these targets, Delete slightly outperformed DiffDec in both mean normalized affinity (MNA Score: 0.472 vs. 0.462) and total number of hits (77.0 vs. 74.3). Furthermore, Delete successfully identified active compounds in 44.3 series, substantially more than the 34.7 series covered by DiffDec. In contrast, DiffDec achieved a higher average number of hits per series (2.15 compared to 1.74 for Delete). As illustrated in **Fig. 6e**, DiffDec demonstrated markedly superior performance over all other methods in both the quantity and potency of active molecules generated for proteins not in CrossDock. This was further supported by the results in **Fig. 6f**, where DiffDec identified active compounds across 18.0 distinct series, exceeding the performance of Delete (13.3), DiffSBDD-C (16.0), and DiffSBDD-M (17.3). Together, these findings underscore the molecular optimization capability of DiffDec in challenging generalization scenarios involving novel proteins, highlighting its robustness and practical utility in structure-based drug discovery. As detailed in the **SI-Fig.9**, our analysis of active molecules across NA score thresholds reveals that DiffDec identifies a greater number of high-affinity ligands for unseen proteins, whereas Delete excels on proteins present during training. This pronounced performance dichotomy underscores the critical necessity of evaluating models on novel protein targets to properly assess their generalizability. Further details are provided in the **Supplementary Information**.

In summary, this comprehensive evaluation underscores the critical role of model architecture, training data composition, and prior knowledge integration in H2L models. While fragment-based models, particularly Delete and DiffDec, demonstrate leading performance across multiple metrics, their strengths are complementary: Although DiffDec excels in discovering active compounds and generalizing to novel targets, its practical application is limited by the need to specify growth sites, an inductive bias that restricts its use when optimal sites are unknown. By contrast, Delete provides broader coverage across diverse molecular series. The superior performance of fragment-based methods over ligand-based approaches reaffirms the value of explicit pocket and fragment information. Our evaluation indicates that current models possess only modest H2L optimization capabilities. Their performance was limited, with the most successful model (Delete) achieving a success rate of just 9.6% (57/600 series). This result highlights a critical limitation and indicates considerable scope for future development. Furthermore, the integration of multi-source data, such as crystallographic structures, simulated complexes, and pharmacophore priors, proves essential for enhancing model robustness and effectiveness. These insights not only illuminate current model capabilities and limitations but also provide a clear pathway for future developments in generative models for drug discovery, emphasizing the need for task-aware design and generalized optimization strategies.

## CONCLUSION

In this study, we first identified critical limitations in the current evaluation of molecular generative models, including narrow dataset scope, underrepresentation of H2L optimization evaluations, lack of metrics to directly evaluate the efficiency and capability of identifying active compounds, lack of generalization testing, and reliance on error-prone computational proxies. We addressed these gaps by curating the most comprehensive, experimentally validated dataset to date for SBDD: this resource comprises 220,005 active molecules across 120 diverse protein targets, paired with 5,433 chemical series to enable rigorous assessment of both de novo generation and H2L optimization. Building on this foundation, we established the first dedicated benchmark for H2L optimization in SBDD, featuring 600 congeneric compound series spanning the same 120 targets, filling a longstanding void in evaluating this critical stage of drug discovery. Using this dataset, we systematically assessed 10 de novo design methods and 7 H2L optimization approaches, spanning diverse architectural paradigms (autoregressive, diffusion, BFN), training data sources (simulated complexes like CrossDock, crystal-derived data like BindingMOAD), and integration of domain-specific prior knowledge (e.g., pharmacophore constraints, protein surface features, chemical rules). This design allowed us to disentangle how each methodological factor shapes performance across SBDD tasks, moving beyond mere method comparison to derive actionable insights.

To probe the generalization potential of these methods in real-world settings, we divided the test proteins into two groups based on their presence in the widely used CrossDock training set, enabling a direct comparison of performance on seen versus unseen proteins. This stratification enabled direct quantification of how prior target exposure inflates performance, a critical oversight in existing benchmarks. Our evaluation framework encompassed a multi-dimensional suite of metrics tailored to SBDD needs: chemical validity, active molecule/scaffold recovery, screening efficiency, coverage of target-specific bioactive space, target-aware capability, conformational quality, and interaction fidelity. For H2L optimization, we further introduced the MNA Score, a novel metric to quantify models’ ability to generate molecules with better binding potency.

Through extensive experimentation, we uncovered consistent limitations across current generative models: both de novo and H2L methods exhibit limited virtual screening efficacy, struggling to generate fully active molecules (even as active scaffold generation remains more feasible). These models also fail to adequately cover target-specific bioactive space in terms of both molecules and scaffolds. Additionally, a significant proportion of generated molecules exhibited high-risk structural motifs (e.g., reactive groups) that would preclude experimental advancement. While increasing sample size could help some methods discover more actives, the absence of reliable scoring functions hinders effective identification of these molecules, limiting practical utility. Another critical finding was the weak target awareness of most structure-based de novo models: despite their design intent to leverage protein structural information, they frequently generate structurally similar molecules across distinct targets, raising concerns about off-target effects in real-world applications. Encouragingly, progress was observed in 3D conformational generation: some methods (e.g., MolCraft for de novo design, DiffDec for H2L optimization) outperformed conventional tools like Vina in specific domains, such as minimizing steric clashes or replicating conserved ligand–protein interactions, which represents a positive development. However, we also noted that common metrics like redocking r.m.s.d. fail to fully capture conformational quality, as they can be inflated by models generating overly rigid molecules, underscoring the need for complementary ligand flexibility analyses (e.g., rotatable bond and ring count). Most importantly, our results highlight whether a protein was seen during training significantly affects performance, suggesting that undifferentiated evaluation risks overestimating practical utility and emphasizing the necessity of rigorous assessment on unseen proteins.

Collectively, these findings demonstrate that substantial advances are still needed before molecular generative models can deliver on their promise of efficient, target-specific molecular design. Our study provides a comprehensive reference that extends beyond mere methodological comparison to offer clear guidance for future development. We hope this work can help build a communicative bridge between AI and life sciences, serving as a foundational resource and practical roadmap to refine generative architectures, improve training paradigms, and integrate more biologically relevant prior knowledge in biological generative AI. Ultimately, this effort aims to accelerate the development of generative models with enhanced clinical impact, speeding the discovery of safe, effective therapeutics for unmet medical needs and realizing the potential of AI to transform SBDD.

## METHODS

### Maximum common substructures matching

Maximum common substructures (MCS) were identified for each chemical series using RDKit. To ensure both representativeness and coverage, we applied a dual filtering criterion: (i) the MCS had to be present in at least 80% of the molecules in the series, and (ii) the substructure had to cover more than one-third of the heavy atoms in each molecule containing it. To satisfy these conditions, the MCS matching threshold was iteratively adjusted from 1.0 downward in steps of 0.1 until both coverage and frequency requirements were met. For each chemical series, the resulting scaffold and the set of molecules meeting the coverage criterion were recorded for subsequent analysis.

### Selection of initial structures and conformations for compound optimization

After confirming the ligands and the PDB structures to be used, we then used Glide^29^ to generate 3D conformations. Initially, the ligands were preprocessed using the Schrödinger LigPrep module with default parameters. On the protein side, the PDB files were prepared using the Protein Preparation Wizard in the Schrödinger suite, following the standard protocol. Water molecules that formed more than three hydrogen bonds with ligand or receptor atoms were retained, and the receptor grids were generated with the centroid of the co-crystallized ligand as the center. After preprocessing, docking was performed using the Glide module in Schrödinger with default parameters. For each resulting ligand pose, the r.m.s.d. of the maximum common substructure relative to all other poses was calculated. The pose exhibiting the lowest mean r.m.s.d. value was selected as the reference conformation for subsequent optimization.

### Molecule generation

In this study, we obtained the selected molecule generation models from their official GitHub repositories. All models were executed using their default configuration settings as specified in the corresponding documentation. For the de novo molecule generation task, each model was required to generate 1,000 molecules per UniProt ID. For the lead optimization task, each model was required to generate 200 molecules per series. To ensure reproducibility and account for stochasticity, each experiment was repeated three times with different random seeds. We provide comprehensive descriptions of the codebases, weight files, input formats, and key parameters for all methods in the **Supplementary Information** file.

To investigate the impact of increased sampling on the efficiency of discovering active molecules for de novo design models, we selected three representative targets based on a specific criterion: the average number of models per target that successfully sampled positive molecules across three independent replicates. According to this ranking, we chose the top two targets from the ‘proteins in CrossDock’ subset and the top one from the ‘proteins not in CrossDock’ subset. For H2L models, we first filtered analog series by requiring r.m.s.d. of maximum common substructure less than 2 Å. These series were then ranked by the number of reference molecules they contained. Following this order, we selected the top two series from the ‘proteins in CrossDock’ subset and the top one from the ‘proteins not in CrossDock’ subset. For these selected proteins or series, we sampled up to 100,000 molecules per protein or series to assess whether increased sampling could enhance the discovery of active scaffolds/molecules and improve the corresponding hit rates.

### Molecule docking

For the Vina docking workflow, ligands were first processed with RDKit to generate initial 3D conformations using the MMFF94 force field and subsequently converted to the PDBQT format using AutoDockTools. The protein structures prepared with the Schrödinger Protein Preparation Wizard directly converted to the PDBQT format for docking in Vina. As commonly practiced in SBDD, the docking grid box was centered on the centroid of the co-crystallized ligand. The box dimensions were determined from the spatial extent of the ligand coordinates, with an additional buffer of 5 Å in each dimension to ensure complete coverage of the binding pocket. Docking simulations were carried out using Vina with default parameters, and the top-ranked binding poses were retained for subsequent analysis. The assessment of docking pose dependent metrics was limited to molecules possessing fewer than 15 rotatable bonds to ensure data quality.

### Evaluation metrics

#### Basic properties

All metrics in this category are normalized to the range [0,1], where higher values indicate better properties. Specifically:

1. Validity evaluates whether a generated molecule corresponds to a chemically valid structure. A molecule is considered valid if it can be successfully parsed and recognized as a complete molecule by RDKit.
2. QED (Quantitative Estimation of Drug-likeness) measures the drug-like properties of molecules using RDKit.
3. Synthetic Accessibility (SA) score estimates the synthetic feasibility of molecules, computed from fragment contributions using RDKit.
4. Uniqueness measures the proportion of unique molecules generated by each model for a given condition.
5. Diversity quantifies chemical diversity among the valid molecules for each target protein:

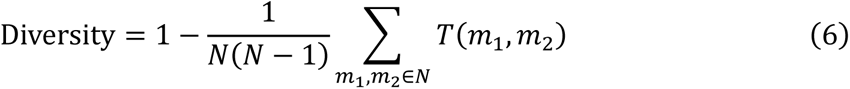

where 𝑇(𝑚_1_, 𝑚_2_) denotes the Tanimoto similarity between molecules 𝑚_1_ and 𝑚_2_, and 𝑁 is the number of valid generated molecules generated for the given target.
6. Chemical similarity to reference actives was evaluated by comparing the distributions of three structural features including Atom types, Ring types, and Functional Group types identified by EFG^55^, between generated molecules and reference molecules. For each feature, empirical distributions were computed and compared using Jensen–Shannon Divergence (JSD). The final score was defined as:

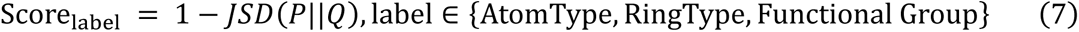

such that higher values indicate closer agreement between generated and reference distributions.

All evaluation metrics except for Validity are computed on molecules that pass our definition of Validity.

### Chemical Filters

To filter out generated molecules with unstable, reactive, or other high-risk properties, we employed the open-source library medchem (https://github.com/datamol-io/medchem). This tool provides a consolidated implementation of authoritative chemical filtering rulesets, integrating proprietary knowledge from industry leaders (such as Eli Lilly’s reactivity rules and Novartis Institute for BioMedical Research’s criteria) with community-driven alerts from public sources like ChEMBL and RDKit catalogs. This offers a robust framework for applying structural filters to prioritize compounds and mitigate risks associated with promiscuity and reactivity, thereby enhancing the efficiency of virtual screening. Details for each chemical filter item in **SI-Table1**.

### Structural plausibility

Structural plausibility of the generated protein–ligand binding poses was evaluated using PoseCheck^35^ and PoseBusters^56^. Using PoseCheck, we computed (i) strain energy to assess the rationality of ligand conformations and (ii) clash score to quantify steric overlaps between protein and ligand atoms. In parallel, PoseBusters pass rates provide a comprehensive evaluation of the physical plausibility of the generated molecular structures, encompassing both intramolecular validity (e.g., internal strain, bond lengths) and intermolecular validity (e.g., steric clashes with the protein). Details for each PoseBuster test item in **SI-Table2**.

### Binding mode consistency

Binding mode consistency of the generated poses was evaluated using two complementary metrics.

### Redocking r.m.s.d

First, to assess binding pose consistency, each generated molecule was re-docked into the corresponding protein pocket using Vina. The r.m.s.d. between the generated pose and the re-docked pose was computed with the spyrmsd^57^ package. Lower r.m.s.d. values indicate higher consistency between the generated and docking-predicted poses.

### Interaction Score

Second, for each target, all reference active molecules were docked to the protein binding site, and interaction fingerprints were computed using ProLIF. Each residue-interaction feature was defined as 𝑘 = (𝑟, 𝑡), where 𝑟 denotes a protein residue and 𝑡 denotes the type of interaction (e.g., hydrogen bond, π–stacking, salt bridge). Residue–interaction features were extracted from all docked reference actives, and the occurrence count 𝑐_𝑘_ of each feature was recorded. We then computed the occurrence rate of each feature and removed those with frequency below a predefined threshold (0.1 in this study). Let 𝒦 denote the set of retained features. The normalized importance weight of each feature was then defined as:

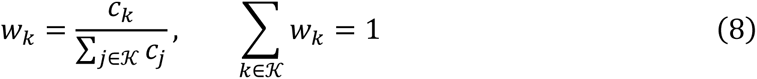

For a generated molecule 𝑔, the set of features it forms is denoted 𝐼(𝑔) ⊆ 𝒦. Its interaction score is defined as:

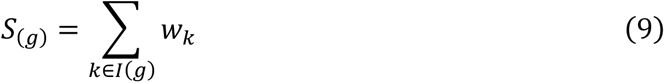

## Acknowledgements

We gratefully acknowledge financial support from National Natural Science Foundation of China (T2225002 and 82273855), Fundamental Research Funds for the Central Universities, National Key Research and Development Program of China (2022YFC3400504 and 2023YFC2305904), the Lingang Laboratory (LGL-8888-02) and Strategic Priority Research Program of the Chinese Academy of sciences (XDB0830200 and XDB1260301).

## Author Contributions Statement

D.C., Z.F. and J.Y. contributed equally. M.Z. and D.C conceived the research project. D.C. and Z.F. developed the primary method and code. J.Y. assisted in the analysis of the primary baselines and data. All authors contributed to the analysis of the results. D.C., Z.F., J.Y. and M.Z. wrote the paper. All authors read and approved the manuscript.

## Data Availability

The data presented in this study are derived exclusively from publicly available datasets. Processed data and all underlying source data for the reported results is available in Zenodo with the identifier https://zenodo.org/records/17572553.

## Code Availability

The code that supports the findings of this study is available in the https://github.com/CAODH/MolGenBench repository under an MIT License.

---

**Extended Fig. 1.**
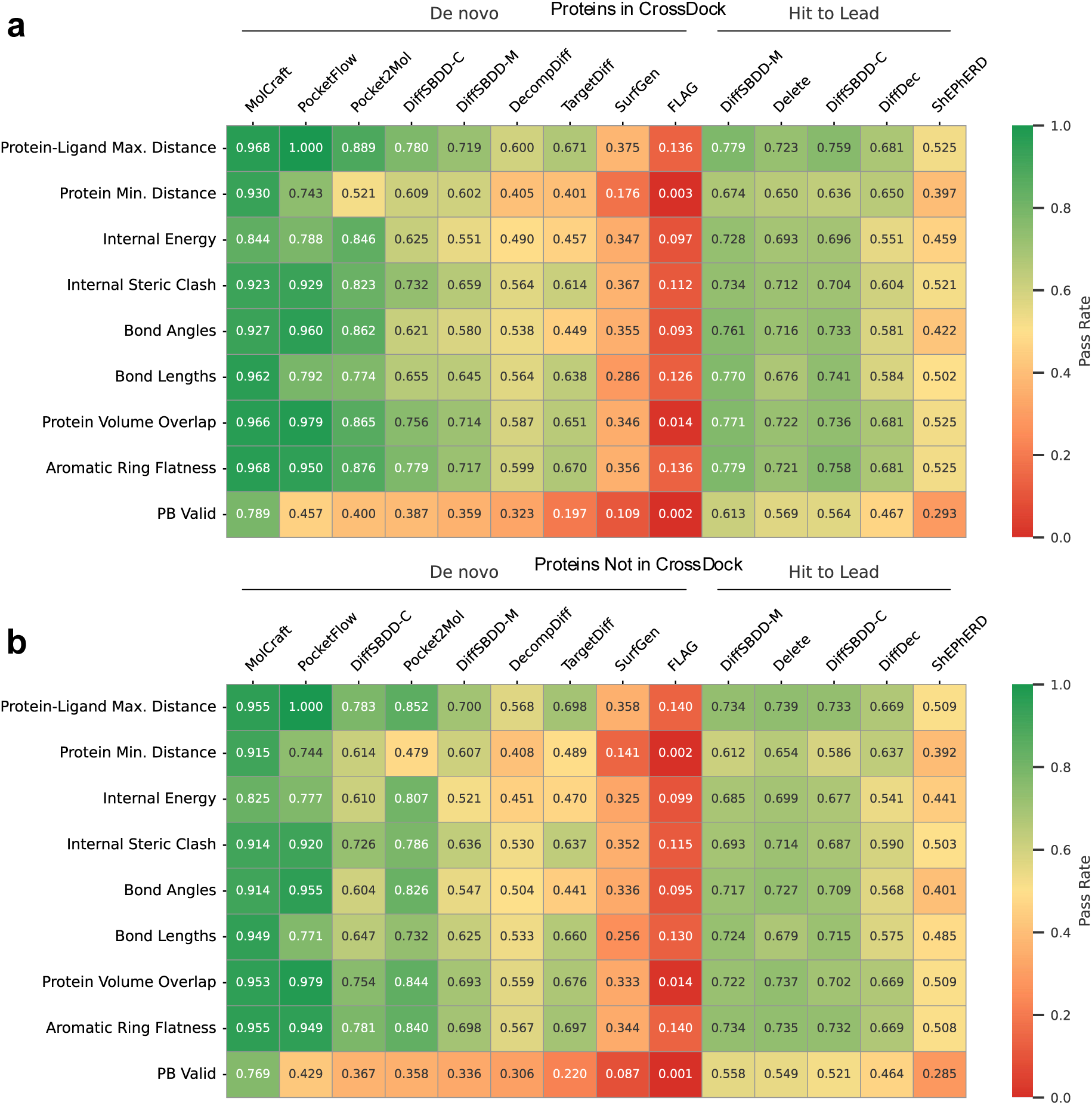
Comparative Evaluation of Conformational Quality Across Generative Models via PoseBusters. **a**, PoseBusters validation results for all generative models (de novo and H2L) on proteins included in the CrossDock training set. **b**, PoseBusters validation results for all generative models (de novo and H2L) on proteins not included in the CrossDock training set. Notably, we calculated the PoseBusters pass rate using the total number of sampled molecules (not just chemically valid molecules) as the denominator. This design inherently classifies invalid or unsuccessfully sampled molecules as conformational failures, providing a holistic measure of performance from initial generation to conformational validation.

**Extended Fig. 2.**
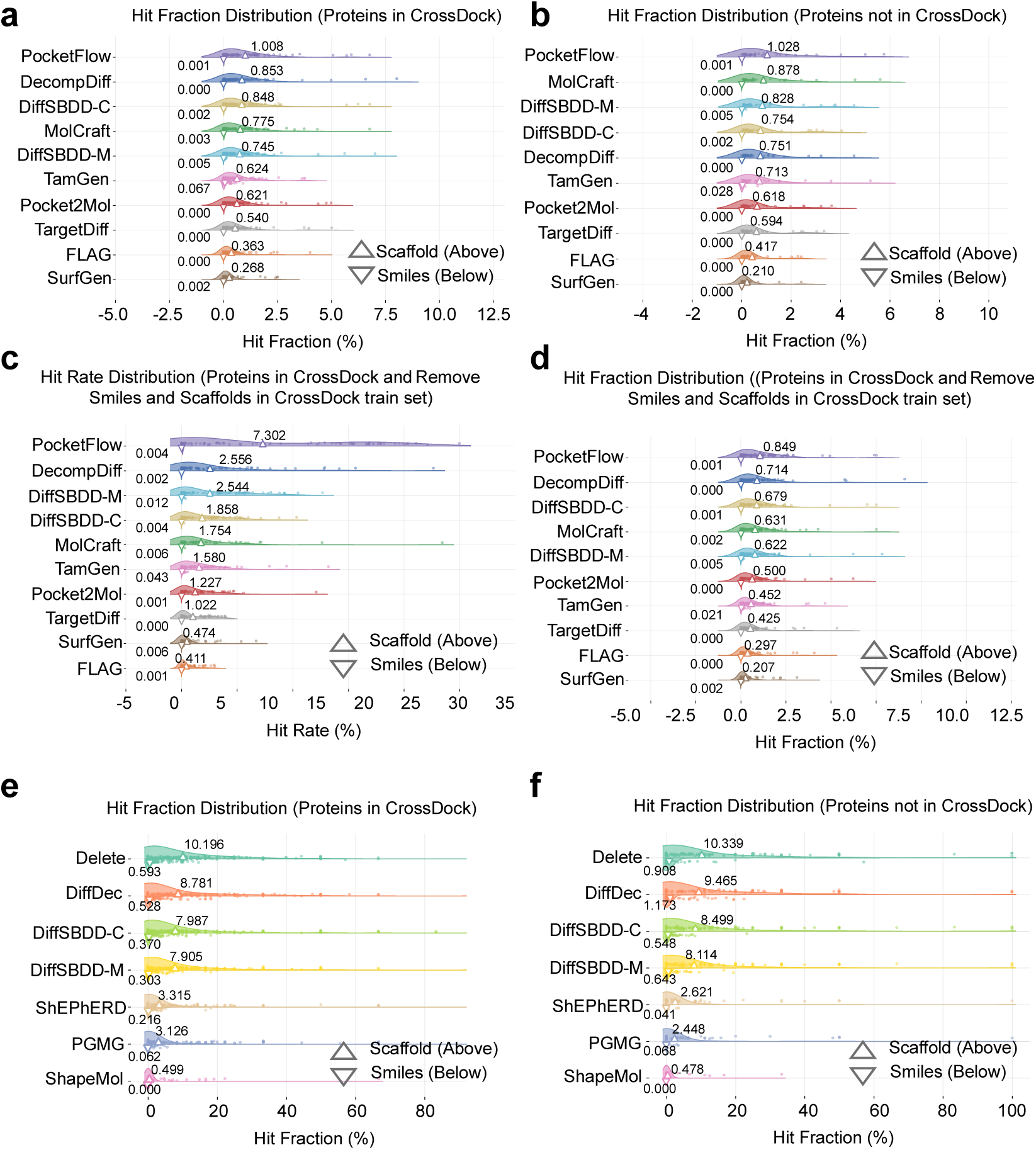
Hit Fraction Evaluation of Molecular Generative Models and the Impact of Training Set Overlap with Target Proteins. **a**, Hit fraction distribution of de novo generative models for proteins included in the CrossDock training set. **b**, Hit fraction distribution of de novo generative models for proteins not included in the CrossDock training set. **c**, Hit rate distribution of de novo generative models for proteins included in the CrossDock training set, after excluding de novo-generated molecules or scaffolds that overlap with the CrossDock training set. **d**, Hit fraction distribution of de novo generative models for proteins included in the CrossDock training set, after excluding de novo-generated molecules or scaffolds that overlap with the CrossDock training set. **e**, Hit fraction distribution of H2L optimization models for proteins included in the CrossDock training set. **f**, Hit fraction distribution of H2L optimization models for proteins not included in the CrossDock training set. Notably, the H2L model DiffDec requires predefined atomic anchors to identify modification sites. Notably, the H2L model DiffDec requires predefined atomic anchors to identify modification sites. These anchors are derived by aligning the common substructure of the compound series to reference actives; shared modification sites across reference actives are weighted by their occurrence frequency, and molecule generation is guided by these sites.

**Extended Fig. 3.**
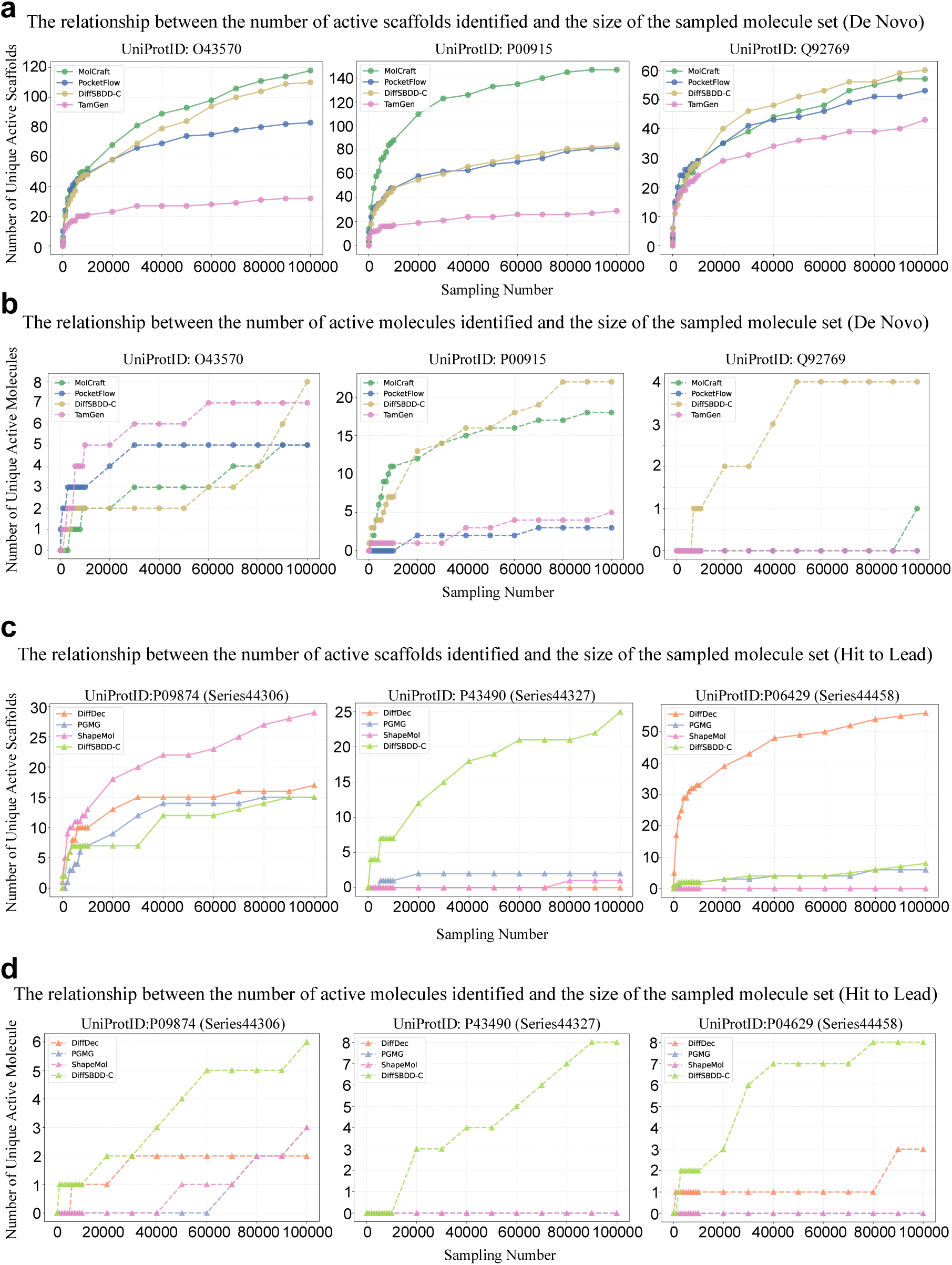
Relationship Between Inference-Time Sampling Volume and Recovery Yield of Active Scaffolds/Molecules in Molecular Generative Models. **a**, Relationship between sampling volume and the recovery yield of active scaffolds in de novo generative models. **b**, Relationship between sampling volume and the recovery yield of active molecules in de novo generative models. **c**, Relationship between sampling volume and the recovery yield of active scaffolds in H2L optimization models. **d**, Relationship between sampling volume and the recovery yield of active molecules in H2L optimization models. Model performance was systematically evaluated across a broad range of sampling volumes, spanning five orders of magnitude (1 to 100,000 molecules per target/series). Intermediate sampling was conducted at 1, 10, 100, and 1,000 molecules, followed by 1,000-molecule intervals up to 10,000 molecules, and 10,000-molecule intervals thereafter to the maximum volume of 100,000 molecules. Notably, the H2L model DiffDec requires predefined atomic anchors to identify modification sites. These anchors are derived by aligning the common substructure of the compound series to reference actives; shared modification sites across reference actives are weighted by their occurrence frequency, and molecule generation is guided by these sites.

