## Supplemental Information File for "Benchmarking Real-World Applicability of Molecular Generative Models from De novo Design to Lead Optimization with MolGenBench"

### Supplementary Information

**Table of Contents**

**S1 Supplementary Information of MolGenBench**

**S2 Supplementary Information for Standard Metrics, Chemical Safety and Scaffold Diversity Evaluation Results**

**S3 Supplementary Information for Conformation Evaluation Results**

**S4 Supplementary Information for Active Molecules Rediscover Evaluation Results**

**S5 Supplementary Information for H2L Evaluation Results**

**S6 Parameters and Inputs of the Molecular Generation Methods Evaluated**

### Supplementary Information of MolGenBench

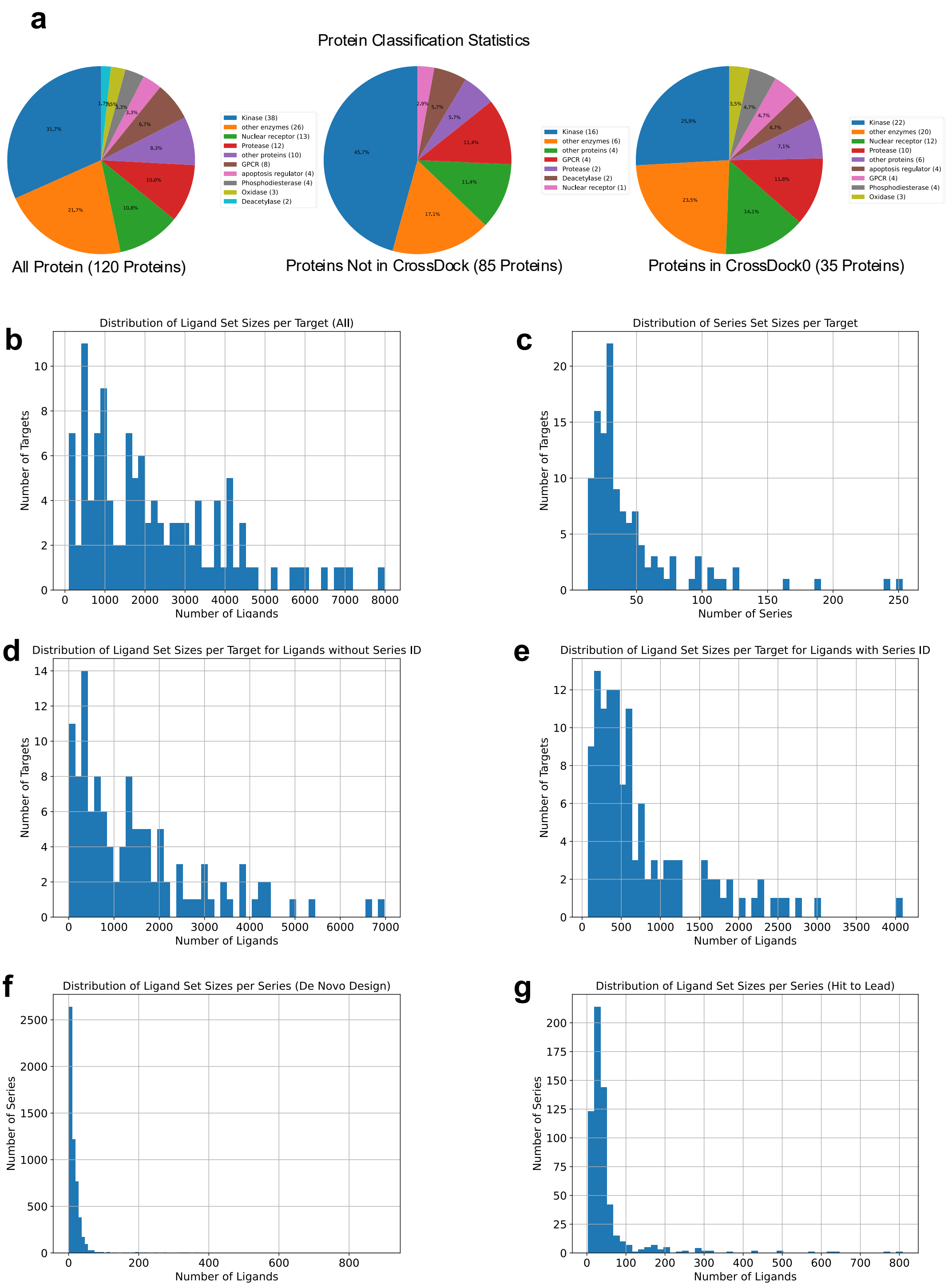


**SI-Fig. 1 | Overview of MolGenBench. a**, Protein classification statistics. **b**, Distribution of ligand set size per target. **c**, Distribution of series set size per target. **d**, Distribution of ligand set size per target for ligand without Series

ID. **e**, Distribution of ligand set size per target for ligand with Series ID. **f**, Distribution of ligand set size per series (De novo Design). **g**, Distribution of ligand set size per series (Hit to lead).

#### Supplementary Information for Standard Metrics, Chemical Safety and Scaffold Diversity Evaluation Results

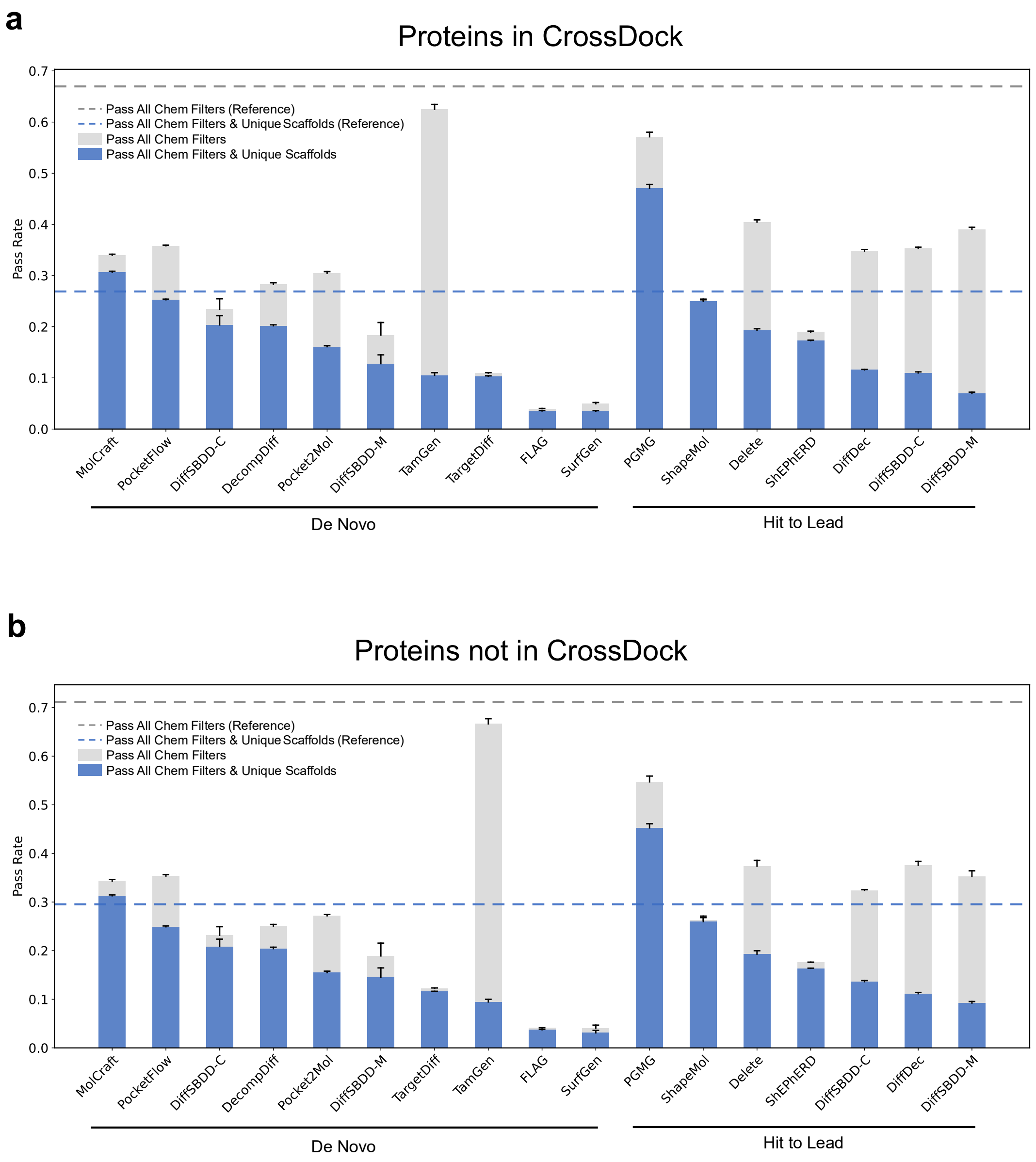


**SI-Fig. 2 | An overview of chemical instability and scaffold diversity in de novo design and molecular optimization across protein subsets. a**, **b**: Chemical stability and scaffold diversity of de novo and molecular optimization models across protein subsets. (**a**) Proteins in CrossDock. (**b**) Proteins not in CrossDock. Bars represent the mean ± standard deviation from three independent replicates.

**SI-Table1| Details for each chemical filter item**

| Name | SMARTS | Catalog description |
|---|---|---|
| Glaxo | 55 | Glaxo Wellcome Hard filters |
| Dundee | 105 | University of Dundee NTD Screening Library Filters |
| BMS | 180 | Bristol-Myers Squibb HTS Deck filters |
| PAINS | 481 | PAINS filters |
| SureChEMBL | 166 | SureChEMBL Non-MedChem Friendly SMARTS |
| MLSMR | 116 | NIH MLSMR Excluded Functionality filters (MLSMR) |
| Inpharmatica | 91 | Unwanted fragments derived by Inpharmatica Ltd. |
| LINT | 57 | Pfizer lint filters (lint) |
| Alarm-NMR | 75 | Reactive False Positives in Biochemical Screens (Huth et al. https://doi.org/10.1021/ja0455547) |
| AlphaScreen-Hitters | 6 | Structural filters for compounds that may be alpha screen frequent hitters |
| GST-Hitters | 34 | Structural filters for compounds may prevent GST/GSH interaction during HTS |
| HIS-Hitters | 19 | Structural filters for compounds prevent the binding of the protein His-tag moiety to nickel chelate |
| LuciferaseInhibitor | 3 | Structural filters for compounds that may inhibit luciferase. |
| DNABinder | 78 | Structural filters for compounds that may bind to DNA. |
| Chelator | 55 | Structural filters for compounds that may inhibit metalloproteins (chelator). |
| Frequent-Hitter | 15 | Structural filters for compounds that are frequent hitters. |
| Electrophilic | 119 | Structural filters for compounds that could take part in electrophilic reaction and unselectively bind to proteins |

| Name | SMARTS | Catalog description |
|---|---|---|
| Genotoxic Carcinogenicity | 117 | Structural filters for compounds that may cause carcinogenicity or/and mutagenicity through genotoxic mechanisms (Benigni rules, https://publications.jrc.ec.europa.eu/repository/handle/JRC43157) |
| LD50-Oral | 20 | Structural filters for compounds that may cause acute toxicity during oral administration |
| Non-Genotoxic-Carcinogenicity | 22 | Structural filters for compounds that may cause carcinogenicity or/and mutagenicity through non-genotoxic mechanisms (Benigni rules, https://publications.jrc.ec.europa.eu/repository/handle/JRC43157) |
| Reactive-Unstable-Toxic | 335 | General very reactive/unstable or Toxic compounds |
| Skin | 155 | Skin Sensitization filters (irritable) |
| Toxicophore | 154 | General Toxicophores |

**Supplementary Information for Conformation Evaluation Results**

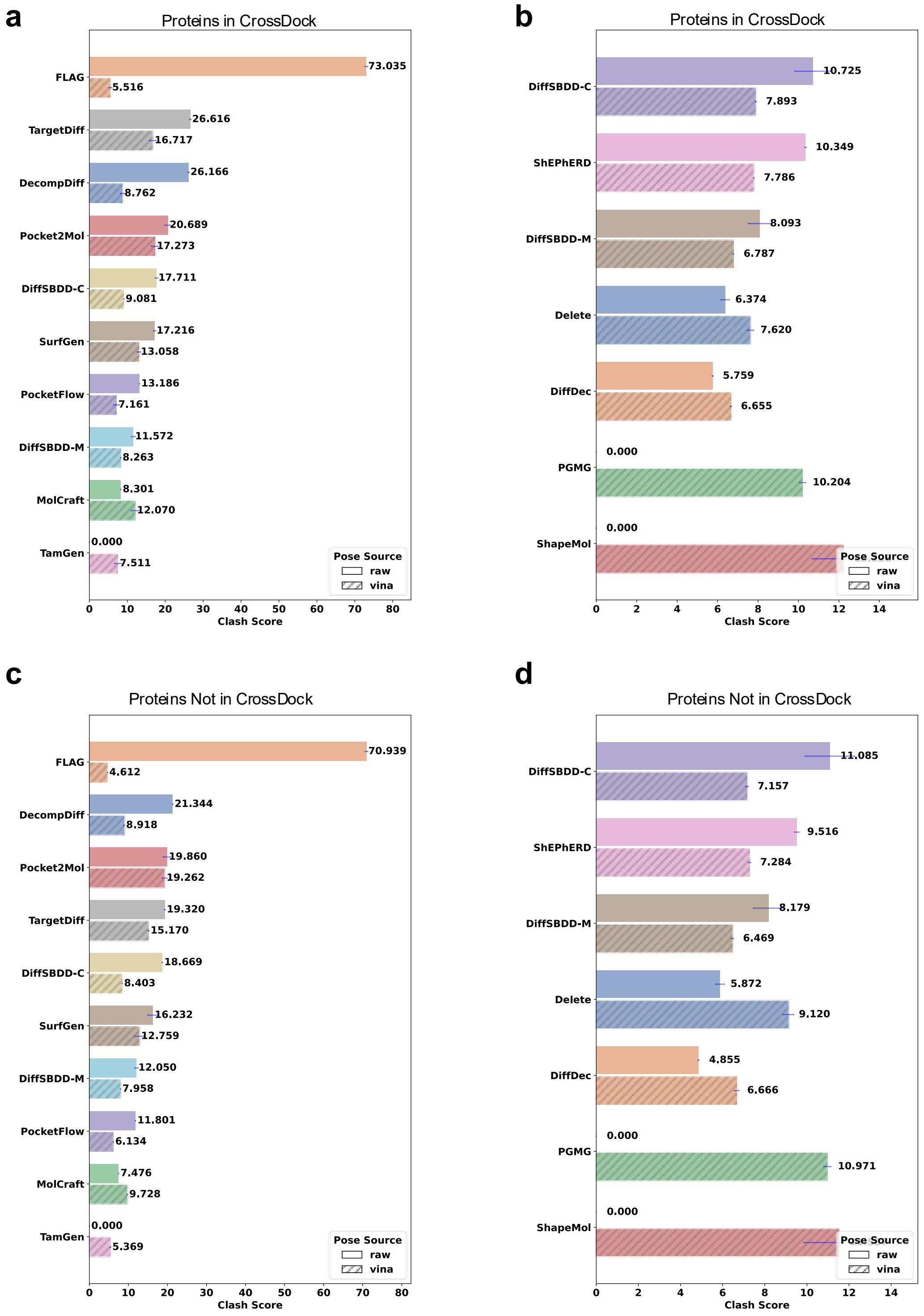


**SI-Fig. 3 | An overview of clash score in de novo design and molecular optimization across protein subsets.** **a**, **c**: Results of clash evaluation for de novo models across protein subsets. (**a**) Proteins in CrossDock. (**c**) Proteins not in CrossDock. **b**, **d**: Results of clash score evaluation for molecular optimization models across protein subsets. (**b**) Proteins in CrossDock. (**d**) Proteins not in CrossDock. Bars represent the mean ± standard deviation from three independent replicates.

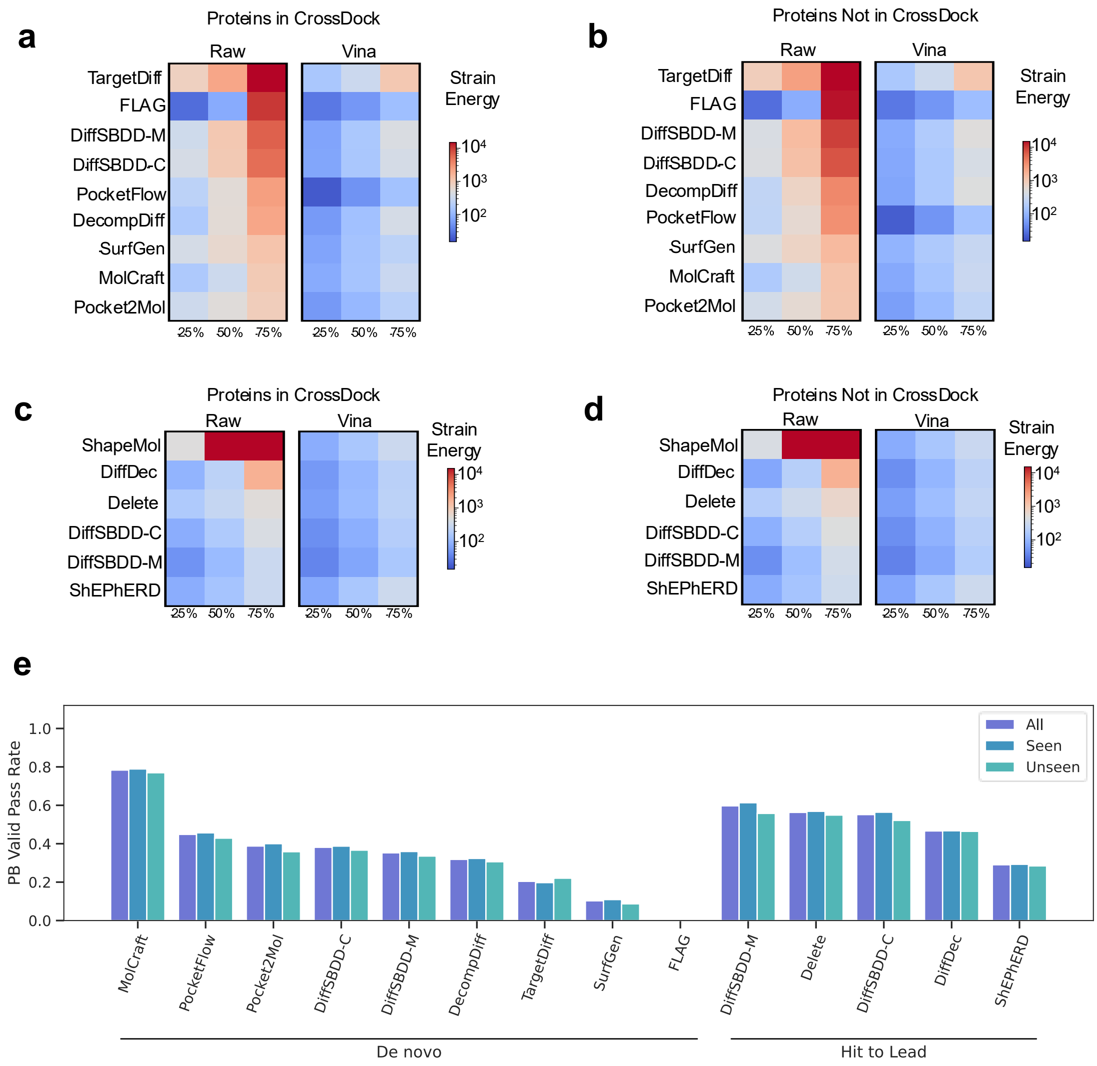


**SI-Fig. 4 | An overview of strain energy and PB-valid pass rate in de novo design and molecular optimization across protein subsets. a**, **b**: Results of strain energy evaluation for de novo models across protein subsets. (**a**) Proteins in CrossDock. (**b**) Proteins not in CrossDock. **c**, **d**: Results of strain energy evaluation for molecular optimization models across protein subsets. (**c**) Proteins in CrossDock. (**d**) Proteins not in CrossDock. Values represent the mean from three independent replicates; **e**, Redocking r.m.s.d. evaluation for all de novo and molecular optimization models on different proteins. Proteins in CrossDock (seen), Proteins not in CrossDock (unseen). Bars represent the mean from three independent replicates.

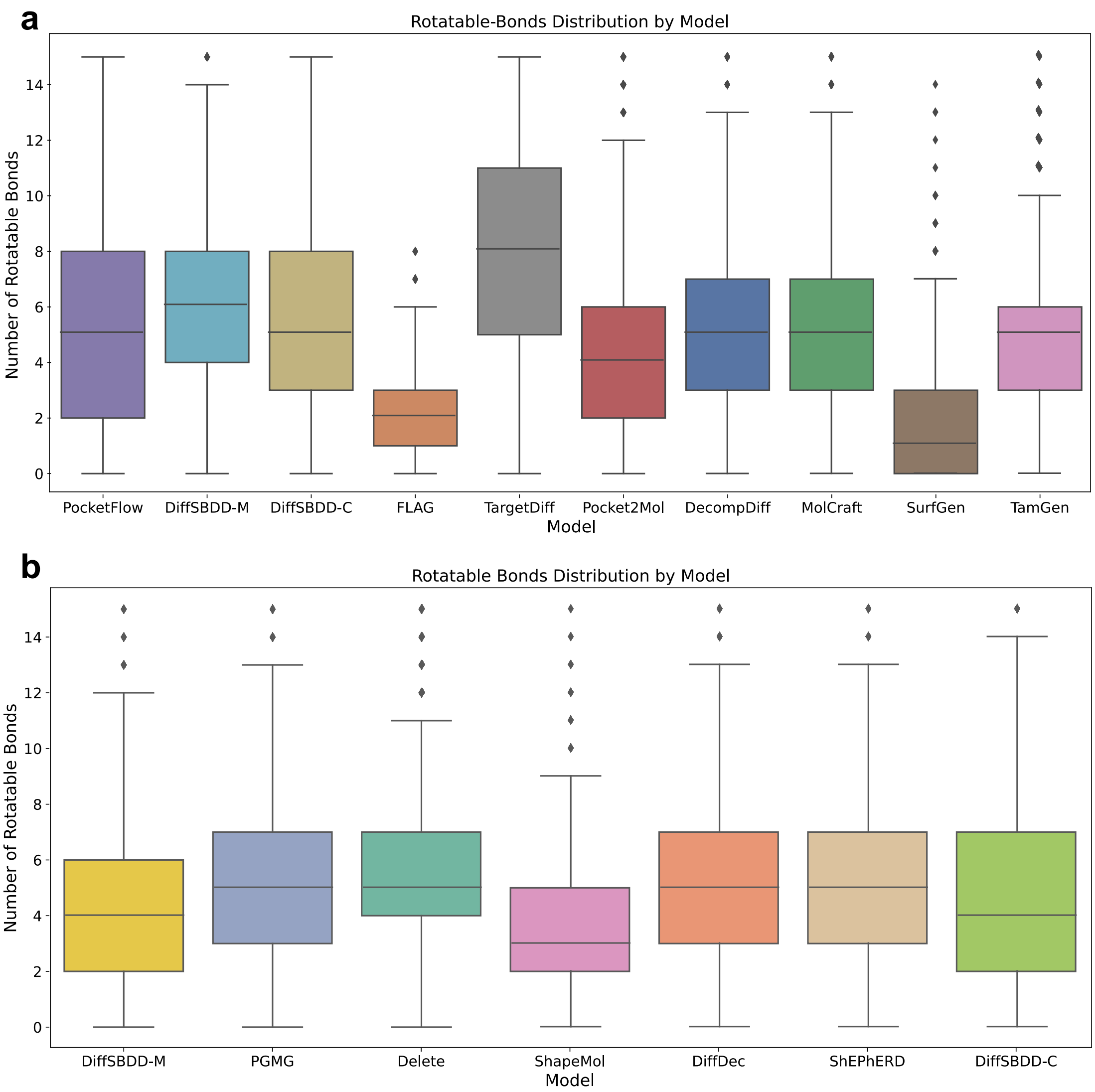


**SI-Fig. 5 | Overview of rotatable bond distribution in de novo design and molecular optimization models.**

**a**, Rotatable bond distribution in de novo design. **b**, Rotatable bond distribution in molecular optimization.

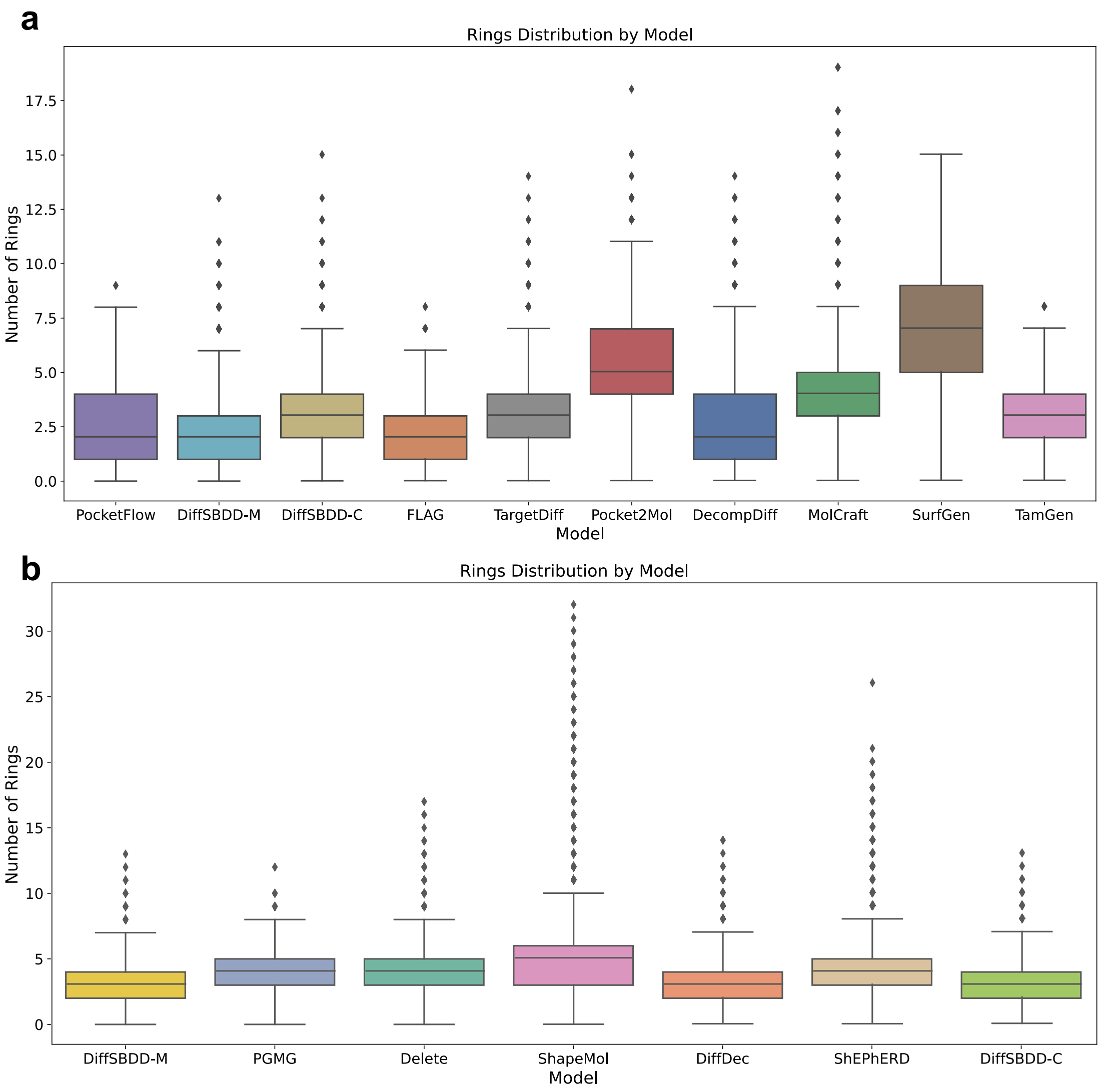


**SI-Fig. 6| Overview of ring distribution in de novo design and molecular optimization models.** a: Ring distribution in de novo design. b: Ring distribution in molecular optimization.

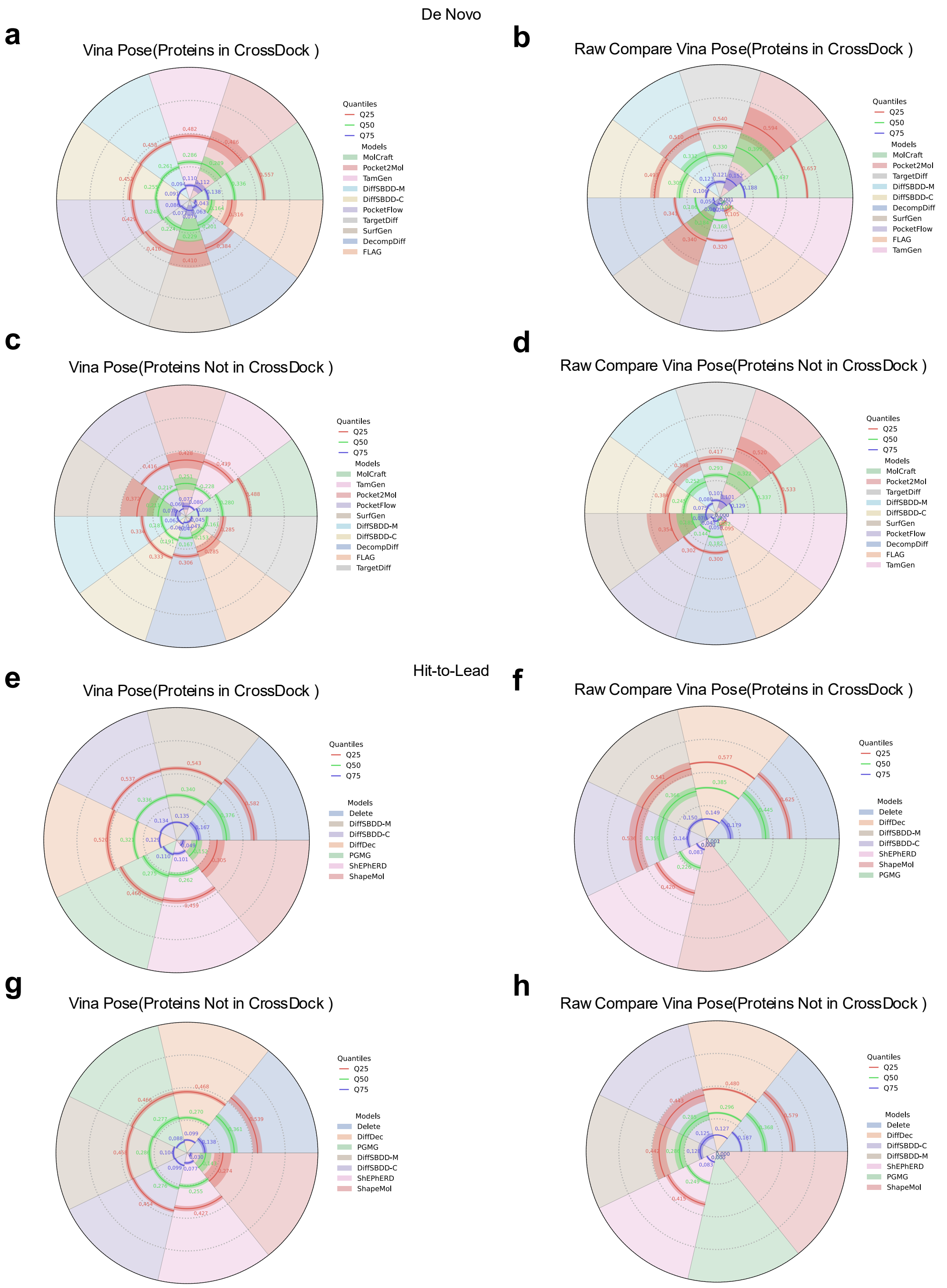


**SI-Fig. 7| Overview of interaction assessment for de novo design and molecular optimization models across protein subsets and ligand pose sources.** Interaction is evaluated for de novo models on proteins (**a**) in and (**c**) not in the CrossDock dataset, using redocked ligand poses. Interaction is evaluated for de novo models on proteins (**b**) in and (**d**) not in the CrossDock dataset, using generated ligand poses. Interaction is evaluated for molecular optimization models on proteins (**e**) in and (**g**) not in the CrossDock dataset, using redocked ligand poses. Interaction is evaluated for molecular optimization models on proteins (**f**) in and (**h**) not in the CrossDock dataset, using generated ligand poses.

**SI-Table2| Specifications for each PoseBuster test item.**

| Property | Description |
|---|---|
| Bond Lengths | The bond lengths in the input molecule are within 0.75 of the lower and 1.25 of the upper bounds determined by distance geometry |
| Bond Angles | The angles in the input molecule are within 0.75 of the lower and 1.25 of the upper bounds determined by distance geometry |
| Protein Volume Overlap | The share of ligand volume that intersects with the protein is less than 7.5%. The volumes are defined by the van der Waals radii around the heavy atoms scaled by 0.8 |
| Aromatic Ring Flatness | All atoms in aromatic rings with 5 or 6 members are within 0.25 A of the closest shared plane |
| Internal Energy | The calculated energy of the input molecule is no more than 100 times the average energy of an ensemble of 50 conformations generated for the input molecule. The energy is calculated using the UFF in RDKit and the conformations are generated with ETKDGv3 followed by force field relaxation using the UFF with up to 200 iterations |
| Internal Steric Clash | The interatomic distance between pairs of non-covalently bound atoms is above 0.7 of the lower bound determined by distance geometry |
| Protein-ligand Max. Distance | The minimum distance between protein-ligand atom pairs is required to be within a 5 Å cutoff. |
| Protein-ligand Min. Distance | The distance between protein-ligand atom pairs is larger than 0.75 times the sum of the pairs van der Waals radii |

#### Supplementary Information for Active Molecules Rediscover Evaluation Results

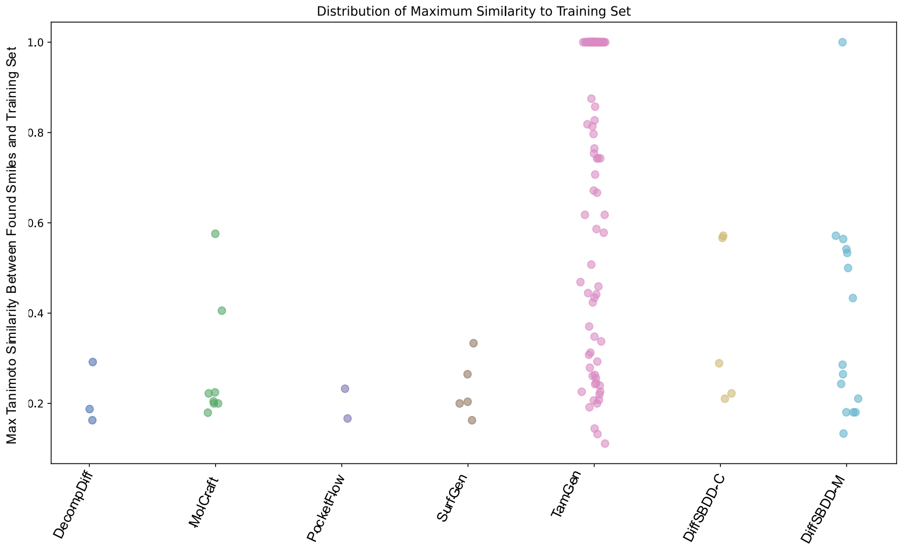


**SI-Fig. 8 | Max Tanimoto Similarity Between Found Smiles and Training Set**

### Supplementary Information for H2L Evaluation Results

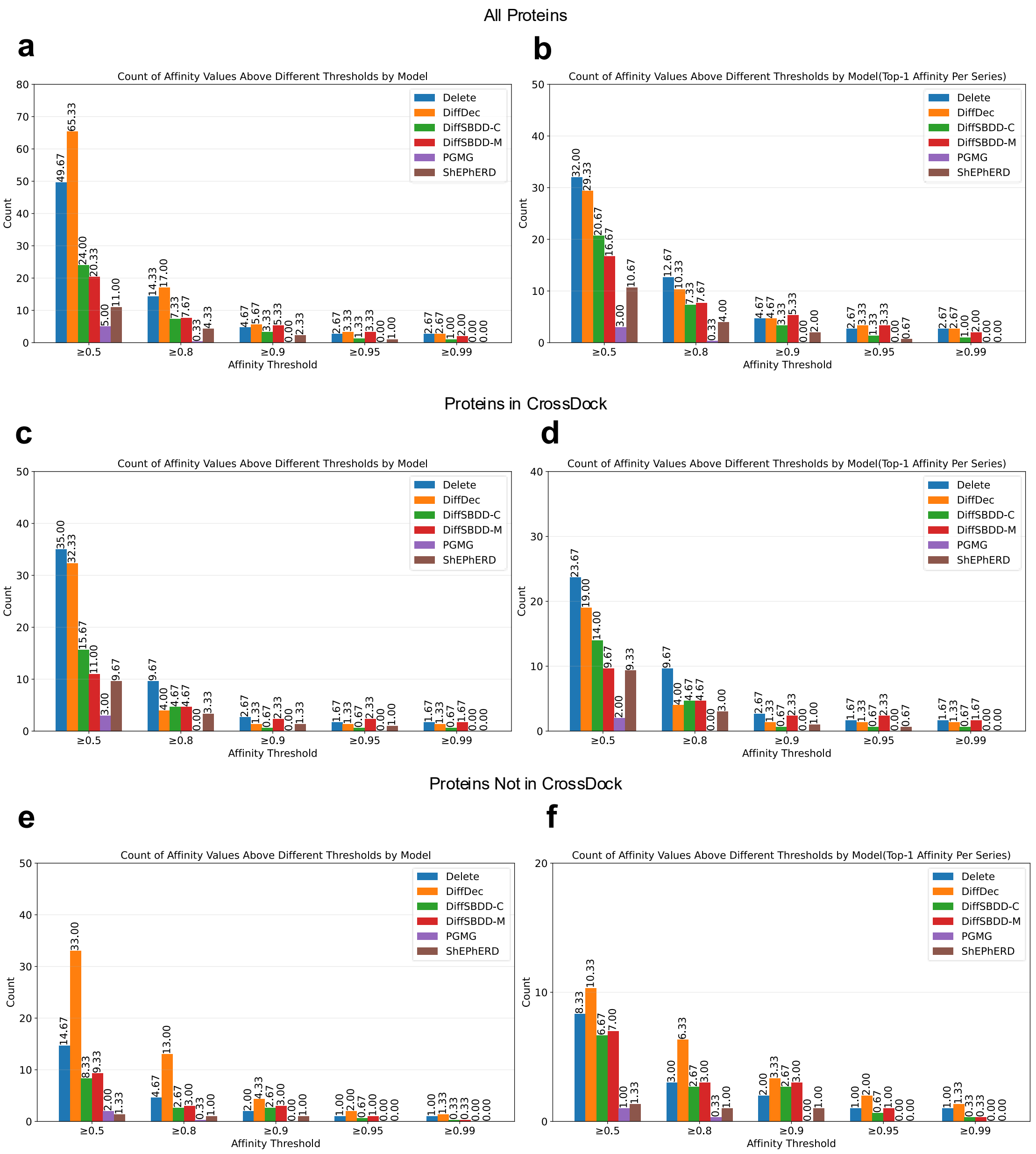


**SI-Fig. 9 | Analysis of active molecules generated by hit-to-lead models across varying thresholds. a,** Number of all active molecules generated by each method across all proteins, evaluated at different thresholds. **b,** For all proteins, the number of active molecules generated by each method when considering only the highest-activity molecule per series across thresholds. **c,** On proteins present in the CrossDock training set, the number of all active molecules generated by each method at different thresholds. **d,** On proteins from the CrossDock training set, the number of active molecules generated by each method when considering only the highest-activity molecule per series across thresholds. **e,** On proteins not encountered during training (unseen in CrossDock), the total number of all active molecules generated by each method at different thresholds. **f,** On unseen proteins, the number of active molecules generated by each method when considering only the highest-activity molecule per series across thresholds. Bars represent the mean value from three independent replicates. Notably, DiffDec requires predefined atomic anchors derived from aligning the common substructure to reference actives to identify modification sites. Shared sites among references were weighted by their occurrence frequency, and molecule generation was distributed accordingly.

We further quantified the number of active molecules identified by each model within their respective series at multiple Normalized Affinity (NA) thresholds: 0.5, 0.8, 0.9, 0.95, and 0.99. As illustrated in **SI-Fig.9a**, DiffDec consistently outperformed all other methods across all protein targets. Specifically, at a threshold of 0.5, DiffDec identified 65.33 active molecules, substantially surpassing Delete, which ranked second with 49.67; other methods yielded markedly lower counts. This performance advantage persisted at higher thresholds, such as at 0.8, where DiffDec again exceeded Delete (17.00 vs. 14.33). Ligand-based methods consistently underperformed relative to fragment-based approaches across all thresholds.

Additionally, we analyzed the maximum NA value per series for molecules identified by each model. At thresholds below 0.8, Delete achieved superior coverage across more compound series. However, at thresholds exceeding 0.9, DiffDec demonstrated performance comparable to, or slightly superior to, that of Delete.

Further stratification by protein type revealed that on targets present in the CrossDock training set, Delete and DiffDec occupied the top two ranks under both evaluation schemes. In contrast, on unseen protein targets excluded from training, DiffDec significantly outperformed all other methods in terms of both total molecule counts and per-series maximum NA values. These findings suggest that Delete performs optimally on familiar protein environments, whereas DiffDec exhibits remarkable generalization capability by identifying more high-affinity molecules for novel targets.

The pronounced performance divergence under different protein settings underscores the necessity of including unseen targets in model evaluation. However, it is important to note that DiffDec requires the specification of growth sites during inference, which introduces a significant inductive bias for generating active molecules. Since the optimal growth site is often unknown in many practical scenarios, this requirement considerably limits its broader applicability.

#### Parameters and Inputs of the Molecular Generation Methods Evaluated

##### De novo design

**MolCraft**

- **Git version:** https://github.com/GenSI-THUAIR/MolCRAFT/tree/282ded4
- **Checkpoint:** *last.ckpt*
- **Input type:** for each UniProtID, the protein pocket was provided as input.
- **Key parameters:** molecules were generated using the *sample_for_pocket.py* script provided in the repository, with *sample_num_atoms* set to prior, and all other parameters set to their default values.

**DecompDiff**

- **Git version:** https://github.com/bytedance/DecompDiff/tree/ed4e7d8
- **Checkpoint:** *uni_o2_bond.pt*
- **Input type:** the protein pockets were processed into datasets using *preprocess_subcomplex.py*, and the resulting datasets were then provided to *sample_diffusion_decomp.py* as inputs.
- **Key parameters:** molecules were generated using the *sample_diffusion_decomp.py* script provided in the repository, with *num_atoms_mode* set to *prior*, *prior_mode* set to *subpocket*, and all other parameters set to their default values.

**FLAG**

- **Git version:** https://github.com/zaixizhang/FLAG/tree/be031f9
- **Checkpoint:** *pretrained.pt*
- **Input type:** following the official repository, the protein pockets were converted into

datasets using *pl.py* and then provided as inputs to *motif_sample.py*.

- **Key parameters:** molecules were generated using the *motif_sample.py* script provided in the repository, with all parameters set to their default values.

**PocketFlow**

- **Git version:** https://github.com/Saoge123/PocketFlow/tree/a31a5a0
- **Checkpoint:** *ZINC-pretrained-255000.ckpt*
- **Input type:** for each UniProtID, the protein pocket was provided as input.
- **Key parameters:** molecules were generated using the *main_generate.py* script provided in the repository, with all parameters set to their default values.

**Pocket2Mol**

- **Git version:** https://github.com/pengxingang/Pocket2Mol/tree/836a0c4
- **Checkpoint:** *pretrained.pt*
- **Input type:** for each UniProtID, the protein pocket was provided as input.
- **Key parameters:** molecules were generated using the *sample_for_pocket.py* script provided in the repository. All parameters were kept at their default values, except that the *center* parameter was set individually for each pocket as the geometric center of its reference active molecule's 3D coordinates.

**SurfGen**

- **Git version:** https://github.com/OdinZhang/SurfGen/tree/9b216de
- **Checkpoint:** *val_119.pt*
- **Input type:** following the instructions in the official repository, each protein pocket was converted into protein surface as input.
- **Key parameters:** molecules were generated using the *gen.py* script provided in the repository, with all parameters set to their default values.

**TargetDiff**

- **Git version:** https://github.com/guanjq/targetdiff/tree/142f1eb
- **Checkpoint:** *pretrained_diffusion.pt*
- **Input type:** for each UniProtID, the protein pocket was provided as input.
- **Key parameters:** molecules were generated using the *sample_for_pocket.py* script

provided in the repository, with all parameters set to their default values.

**TamGen**

- **Git version:** https://github.com/microsoft/TamGen/tree/2b0c948
- **Checkpoint:** *checkpoint_best.pt*
- **Input type:** for each UniProtID, the input dataset was prepared using the *prepare_pdb_ids_center.py* script, in which the binding-site center was defined by the geometric center of the reference active ligand.
- **Key parameters:** molecules were generated by following the *generator-7vh8.sh* script provided in the repository, using the unconditioned generation mode, with all other parameters set to their default values.

**DiffSBDD-C**

- **Git version:** https://github.com/arneschneuing/DiffSBDD/tree/a2c64e6
- **Checkpoint:** *crossdocked_fullatom_cond.ckpt*
- **Input type:** for each UniProtID, the protein pocket was provided as input.
- **Key parameters:** molecules were generated using the *generate_ligands.py* script provided in the repository, with all parameters set to their default values, except that *all_frags* option was enabled to avoid discarding smaller fragments and to allow evaluation of molecular connectivity.

**DiffSBDD-M**

- **Git version:** https://github.com/arneschneuing/DiffSBDD/tree/a2c64e6
- **Checkpoint:** *moad_fullatom_cond.ckpt*
- **Input type:** for each UniProtID, the protein pocket was provided as input.
- **Key parameters:** molecules were generated using the *generate_ligands.py* script provided in the repository, with all parameters set to their default values, except that *all_frags* option was enabled to avoid discarding smaller fragments and to allow evaluation of molecular connectivity.

**Lead optimization**

**PGMG**

- **Git version:** https://github.com/CSUBioGroup/PGMG/tree/85fb712
- **Checkpoint:** *chembl_fold0_epoch32.pth*

- **Input type:** the active reference molecule in each SeriesID was used to extract pharmacophore information, which was stored in the *edgep* format. These processed inputs were then provided to the *generate.py* script.
- **Key parameters:** molecules were generated using the generate.py script provided in the official repository, following the demo usage. All parameters were kept as default, except that the *filter* option was disabled to avoid restricting the outputs to only unique and valid molecules.

**DiffDec**

- **Git version:** https://github.com/biomed-AI/DiffDec/tree/916ae14
- **Checkpoint:** *diffdec_multi.ckpt*
- **Input type:** for each SeriesID, the protein pocket, the binding pose of the common substructure (obtained by splitting from the reference active molecule in the series), and the corresponding scaffold SMILES with annotated modification anchors were provided as inputs. The scaffold SMILES were obtained by matching the extracted common substructure to the active molecules in the same SeriesID to identify plausible modification sites. Because different actives may match the common substructure differently, multiple scaffold SMILES can arise within a single SeriesID; the total of 200 generated molecules was allocated across these scaffold–modification strategies in proportion to the number of actives that matched each scaffold.
- **Key parameters:** molecules were generated using the *sample_multi_for_specific_context.py* script provided in the repository, with all parameters set to their default values.

**ShEPhERD**

- **Git version:** https://github.com/coleygroup/shepherd/tree/21e79b8
- **Checkpoint:** *x1x3x4_diffusion_mosesaq_20240824_submission.ckpt*
- **Input type:** the active reference molecule in each SeriesID was converted into molblocks, and its charges were extracted; these processed inputs were then provided to the *RUNME_conditional_generation_MOSESaq.ipynb* notebook as the required inputs.
- **Key parameters:** molecules were generated by following the instructions provided in the *RUNME_conditional_generation_MOSESaq.ipynb* notebook included in the repository, using the default settings specified therein, with *n_atoms* set to the number of atoms in the reference molecule to ensure an appropriate atom count.

**ShapeMol**

- **Git version:** https://github.com/Amelie-Schreiber/ShapeMol/tree/main
- **Checkpoint:** *MOSES2_test_mol.pkl*
- **Input type:** binding pose of the reference active molecule in the series were provided as inputs.
- **Key parameters:** *Run ~/ShapeMol/source/scripts/sample_diffusion.py by using the parameters in config file ~/ShapeMol/config/sampling/SMG/sample_diff_pos0_10_pos1.e-7_0.01_6_v001_bondTrue_scalar128_vec32_layer8_with_guidance.yml*

**Delete**

- **Git version:** https://github.com/HaotianZhangAI4Science/Delete/tree/52dff5e
- **Checkpoint:** *crossdock_val_159.pt*
- **Input type:** for each SeriesID, the protein surface (obtained by runing *~/utils/massif/generate_prot_ply.py)* binding pose of the common substructure (obtained by splitting from the reference active molecule in the series) were provided as inputs.
- **Key parameters:** In our implementation, the optimization starting point was defined by our maximum common substructure, replacing the scaffold extraction method provided in the original code.

**DiffSBDD-C**

- **Git version:** https://github.com/arneschneuing/DiffSBDD/tree/a2c64e6
- **Checkpoint:** *crossdocked_fullatom_cond.ckpt*
- **Input type:** for each SeriesID, the protein pocket and the binding pose of the common substructure (obtained by splitting from the reference active molecule in the series) were provided as inputs.
- **Key parameters:** molecules were generated using the *inpaint.py* script provided in the repository, with all parameters set to their default values, except that *all_frags* option was enabled to avoid discarding smaller fragments and to allow evaluation of molecular connectivity.

**DiffSBDD-M**

- **Git version:** https://github.com/arneschneuing/DiffSBDD/tree/a2c64e6
- **Checkpoint:** *moad_fullatom_cond.ckpt*
- **Input type:** for each SeriesID, the protein pocket and the binding pose of the common

substructure (obtained by splitting from the reference active molecule in the series) were provided as inputs.

- **Key parameters:** molecules were generated using the *inpaint.py* script provided in the repository, with all parameters set to their default values, except that *all_frags* option was enabled to avoid discarding smaller fragments and to allow evaluation of molecular connectivity.
